# The drug disposition in the Spleen: A novel study of CYP and UGT-mediated drug metabolism

**DOI:** 10.1101/2024.04.17.589706

**Authors:** Patience Inuwa, Stuart Paine, Stuart Best, Karolina Mosinska-Kodzik, Cyril Rauch

## Abstract

Understanding the fate of a drug, its disposition and pharmacokinetics as it reaches the site of action is key to any pharmaceutical research and development. The emerging role of the spleen in its involvement in regulating the immune system has garnered interest in new immunotherapeutic strategies. Using novel precision immunotherapeutic drugs that will potentially engage with the host immune system to specifically target and eliminate diseased cells, makes this approach a better alternative to conventional therapies. The spleen poses as a potential immunotherapeutic target and confirmation of drugs reaching its site of action requires monitoring enzymes that engages with xenobiotics like drugs. The pig was selected as the model for human based on the close homology conveyed between pig and human using a phylogenetic construction of Cytochrome P450s (CYP450) and UDP-glucuronosyltransferases (UGT) in both species. Moreover, an RNAseq transcriptome analysis between the human spleen, pig liver and spleen tissues were obtained from a next generation sequencing (NGS) to identify genes associated with drug metabolism. Immunofluorescence and Western Blot analysis was carried out to determine the protein expression of metabolizing CYP450s and UGTs in pig spleen. Therefore, drug substrates and their metabolites known in human liver were investigated in pig spleen to determine the functional expression of CYP450s and UGTs. Promising *in-vitro* results has demonstrated the expression of these metabolic enzymes at a functional level from observations showing elimination of drug substrates and the apparent metabolites formed. Monitoring the enzyme activities would also indicate uptake of these substrates in splenocytes, confirm that the spleen can metabolize drugs, and provide further insight into therapeutic or toxic related implications from drug interactions.

## Introduction

The spleen is the second largest of the immune organs and has emerged as an area of interest for targeted immuno-oncology therapies. Secondary immune organs are responsible for generating immune responses and tolerance against foreign invaders (Ruddle & Akirav, 2009). With a host of immune cell population and its ability to eradicate bacterial infections, the spleen has the potential to establish anti-viral, anti-bacterial, and anti-tumor immunity in the body. For example, mainstream oncology therapies commonly rely on invasive surgeries, radiotherapy and chemotherapy or combination of such approaches to increase efficacy, but clinical outcomes of these approaches have been unsatisfactory due to severe adverse and toxic effects including low selectivity and specificity (Bush & Noe, 2020; Ruddle & Akirav, 2009). Hence, recent developments have prompted a demand for new strategies particularly in stimulating the immune response. The concept of immunotherapy has revolutionized the treatment paradigm of cancer. It involves stimulating a patient’s own immune system to recognize and eliminate tumor cells (Bush & Noe, 2020; Conn, 2018). Enhancing the key aspect of immunotherapy would increase the activation of the adaptive immune system primarily through effector T cells and cytotoxic T cells (Conn, 2018; W. Li et al., 2020). It is therefore essential to optimize the stimulation of the immune system concentrating on lymphoid organs like the spleen that contains a high proportion of immune cells. In this context the design of successful drug candidates that would be introduced into the spleen, and then stimulate immune cells should have these important considerations: a) the ability to reach the target b) exhibit good distribution into and within the target tissue and, c) show evidence of its presence at the target and engages with receptors that would induce a specific immune response. This shows that a better understanding of the potential role of the spleen in drug metabolism is paramount.

It is understood that the main site for drug metabolism in the human body is the liver organ. This is due to the high expression of drug metabolizing enzymes known as cytochrome P450s (CYP450) and UDP-glucuronosyltransferase enzymes (UGTs) which are involved in Phase I (functionalization) and Phase II (conjugation) reactions, respectively. These enzymes maintain the chemical homeostasis (or the fate of xenobiotics) in the human body (Mackenzie et al., 2017) and are concentrated in microsomes (of the endoplasmic reticulum) for use in the estimation of Human ADME (Absorption, Distribution, Metabolism and Excretion) (Bachmann, 2009; Vrbanac & Slauter, 2016). Drug metabolism represents an elimination mechanism for most drugs and becomes a key determinant for drug design (Y. Li et al., 2019; Mackenzie et al., 2017). While the liver has gained a lot of attention, it is worth recalling however that drug metabolizing enzymes have been found in various tissues such as the lungs, kidneys, liver, and placenta (Caldwell et al., 1995; Zhang & Tang, 2018). Thus, this raises the question about the possibility that other organs, apart from the liver, can also metabolize drugs efficiently.

To achieve a clear understanding of drug metabolism, the most cost-effective way to study drug metabolism relies on an animal model. Pigs are an ideal animal model for human health and disease due to their similarity in anatomy and physiology and with the recent completion of the swine genome sequence (Walters & Prather, 2013). Most systems and organs in pigs like the heart, liver, kidney, brain, nasal cavity, reproductive and gastrointestinal system, show analogies to human with significant advantages over other experimental models, e.g., rodent and primate counterparts (Puccinelli et al., 2011). The use of pigs also allows for safe dosage ranges to be defined in drug development studies and toxicological testing since they have similar responses to several variety of drugs (Lunney et al., 2021; Walters & Prather, 2013). Understanding the mechanisms of diseases is of importance and animal models like pigs can be used to observe for example differences of drug exposure (Ayuso et al., 2021). The importance of the pig in drug metabolism can be simply demonstrated using a phylogenetic tree comparing human and pig genomes, Figure 1 shows examples of homologies in drug metabolizing enzymes. Therefore, pigs can potentially be used to study drug metabolism in different organs like the spleen.

**Figure 1:**
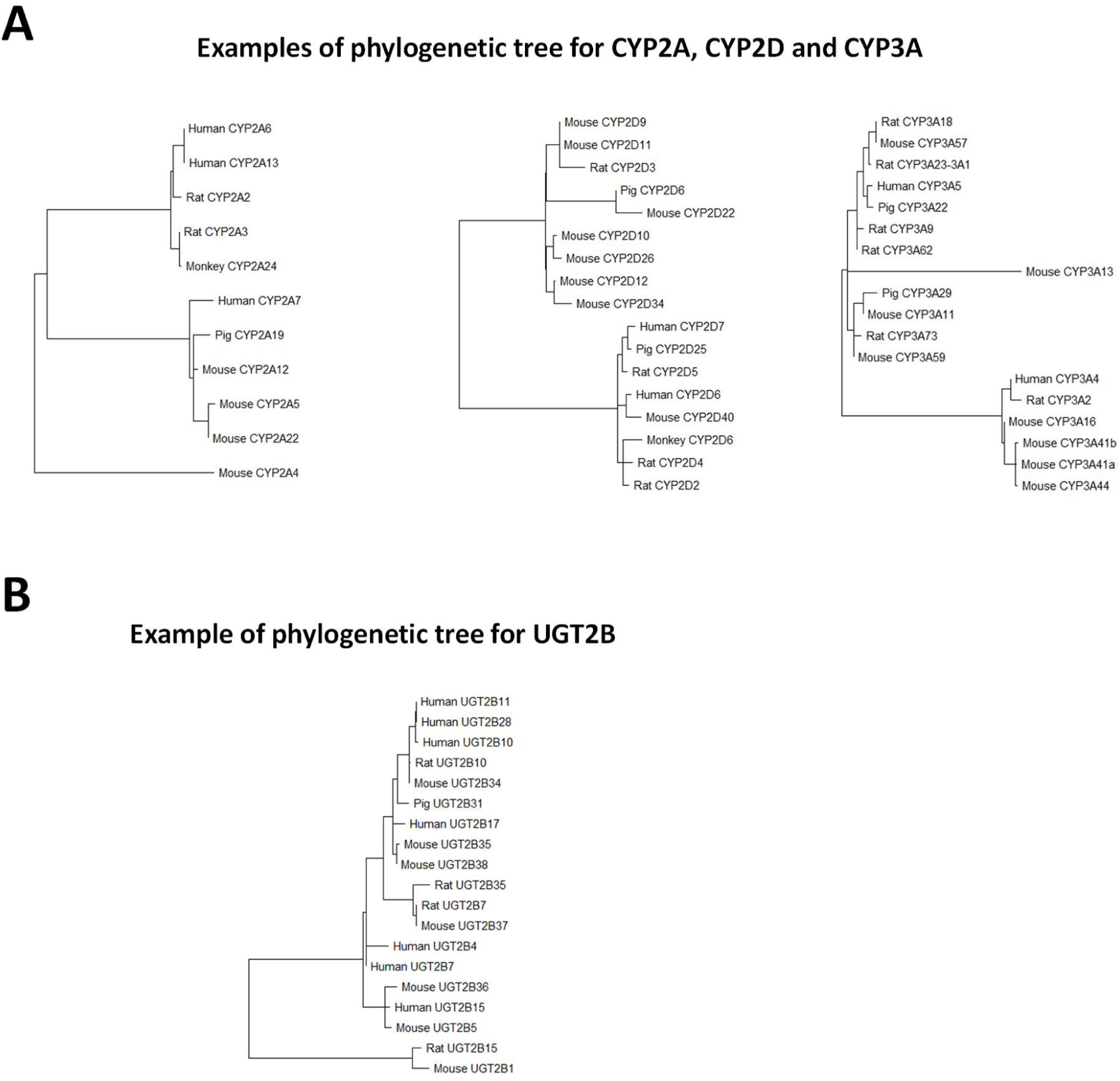
Example of CYP450s and UGTs mRNA sequences obtained from NCBI database.

In summary, the emerging interest in the spleen as a potential therapeutic target together with its potential role in drug metabolism has become significant. This study will focus on the drug disposition in the spleen by investigating the genetic, protein and functional expression of CYP450 and UGT enzymes in the pig spleen.

## Materials and Methods

### Materials and Reagents

Umbelliferone, Serotonin hydrochloride, Naloxone hydrochloride dihydrate, Midazolam solution (in methanol), Omeprazole, Amodiaquine dihydrochloride dihydrate, Astemizole, Dextromethorphan hydrobromide, NADPH (Tetrasodium Salt), UDPGA, Alamethicin, Bovine serum albumin (BSA), dimethyl sulfoxide (DMSO), 2-Mercaptoethanol (β- ME) and Magnesium chloride (MgCl) were purchased from Sigma Aldrich, UK. 10x Phosphate Buffered Saline (PBS) and Laemmli buffer was purchased from Bio-Rad Laboratories, UK. Water (LC-MS grade), Acetonitrile (LC-MS grade), Paraformaldehyde and Formic acid were obtained from Fisher Scientific UK. Liver and spleen tissues stored in dry ice were harvested from female Landrace x Large white x Duroc (domestic pig) post-mortem, in accordance with the procedures approved in schedule 1 of the Animal Welfare Act 2006. The tissues were obtained from the abattoir at Vet school in University of Nottingham and immediately stored in -80° C. The primary antibodies used (*CYP2E1*, *CYP450* generic, *CYP20A1*, *CYP27A1*, *UGT8*) were supplied by Thermo Fisher UK.

### Next Generation Sequencing (NGS)

The NGS analysis using two individual tissue samples of pig liver and spleen was outsourced to Source Bioscience UK. The analysis involved two main steps in the workflow: Sample and Sequencing. The Sample workflow involves quality control and validation of libraries to be sequenced. The Sequencing workflow involves the quantification of gene expression (RNA-seq bioinformatics). The RNA extraction was performed using the Qiagen RNeasy Mini Kit. The tissue was lysed and then subsequently purified in duplicate. Following extraction, the concentration and integrity of the RNA sample were assessed using Invitrogen Qubit RNA assay (Agilent) and BioAnalyzer 2100 (Agilent), respectively. For all RNA samples, the RIN number was above 8. For the library preparation, 500 ng of total RNA was used. The libraries were prepared using the NEBNext Ultra II Directional RNA Library Preparation Kit (NEB), according to the manufacturer’s protocol. During this process, the libraries were indexed using NEBNext Multiplex Oligos for Illumina 96 Unique Dual Index Primer Pairs (NEB). The prepared libraries were quantified via a fluorometric method involving Qubit High Sensitivity (Invitrogen) and qualified using electrophoretic separation on the Bioanalyzer HS Kit (Agilent). The libraries were normalized and pooled before sequencing on the NovaSeq 6000 (Illumina) instrument.

### Protein Lysate preparation

Protein lysates from pig liver and spleen tissues was prepared using a T-PER Tissue Protein Extraction Reagent (#78510, Thermo Fisher, UK) according to the manufacturer’s instructions. Briefly, tissues were homogenized with the reagent, 1 gram of tissue to 10 mL of T-PER Reagent. After centrifugation at 10, 000 xg for 5 min, the pellet tissue debris was discarded, and the supernatant was collected. Total protein concentration was determined using the Bradford method (Nicholas J. Kruger, 2002; Redmile-Gordon et al., 2013).

### Western Blot

Aliquots of the protein lysates from pig and liver tissues (10 µL/ well) were resolved on 4-15 % SDS-polyacrylamide electrophoresis (SDS-PAGE) gel and transferred onto nitrocellulose (NC) blots. The protein extracts were normalized with tissue-lysis buffer (to the desired protein concentration), and an equal volume of Laemmli buffer (consisting of 60 mM Tris-HCl pH 6.8; 20% glycerol; 2% SDS; 4% β-mercaptoethanol; 0.01% bromophenol blue) to a 1:1 volume ratio (Gavini K & Parameshwaran K, 2023). The prepared samples were heated at 95 °C for 10 min. The nitrocellulose blots were blocked using Everyblot buffer (12010020, Bio-Rad UK) for 1 hr at RT, and incubated with primary antibody diluted (1:2000) in Everyblot buffer overnight at 4 ◦C. Then nitrocellulose membranes were washed with TBST and incubated with HRP-conjugated goat anti-rabbit IgG secondary antibody (Thermo Fisher, UK) diluted 1:10000 in Everyblot buffer for 1 hr at RT. The proteins were detected using Clarity Western ECL Substrate (1705061, Bio-Rad UK) chemiluminescence reagent. The chemiluminescence signal was visualized using the ChemiDoc Imaging System (1708370, Bio-Rad UK). Negative control (absence of primary antibodies) was also performed to validate the results obtained.

### Single-cell suspension preparation and isolation of primary cells

Four samples of 50 mg liver and spleen solid fresh tissues were soaked for 15 mins in a dissociation EasySep buffer (20144, Stem Cell Technologies, UK) containing Dulbecco’s phosphate-buffered saline (PBS), 2 % fetal bovine serum (FBS) and 1mM EDTA. The tissues were minced and crushed using a piston/ flat end of plunger to release the cells, which were incubated on rotation at room temperature for 30 min. The cell suspension was filtered and centrifuged at 300 xg for 10 mins. The supernatant was discarded, and the pellet was resuspended in media ready for use within 1 hr of harvesting. The cell viability was above 72 % as determined with the trypan blue exclusion test. Viability was also tested after thawing.

### Immunofluorescence – staining (IF)

The cell suspension (containing 10^6^ cells) from pig liver and spleen tissues were fixed with 4 % paraformaldehyde in PBS, then incubated at room temperature for 20 min. After washing with PBS (2 x 5min) the cells were blocked and permeabilized using 0.05 % Saponin/ 2 % BSA in PBS for 1 hr, the plated cells were incubated with primary antibodies for 1 hr at room temperature. The primary antibodies were diluted in (1:500) in 0.05 % Saponin/ 2 % BSA in PBS and at the antibody concentration recommended by the manufacturers, which were detected with the corresponding goat anti-rabbit secondary antibody conjugated to a fluorochrome (Alexa Fluor™ 488, A-11008, Invitrogen, Thermo Fisher UK). Straight-after, the plated cells were incubated with DAPI for 30 min at room temperature in the dark, then washed with 0.1 % BSA in PBS three times following staining. The cells were mounted on slides and visualized using Leica inverted fluorescent microscope.

### Preparation of microsomes

Microsomes were prepared by differential centrifugation. Liver and spleen tissues (1 g per 10 mL) were homogenized at 4 °C in 0.15 M potassium chloride buffer and centrifuged at 10, 000 xg for 20 min. The supernatant was centrifuged at 100, 000 xg for 1 hr. The supernatant was discarded, and the pellet was suspended in 0.1 M potassium phosphate buffer (pH7.4) and further centrifuged at 100, 000 xg for 1 hr. The microsomal pellet was re-suspended in 0.1M potassium phosphate buffer pH7.4 and stored at -80 °C until use. Protein concentration was determined by the Braford method (Nicholas J. Kruger, 2002; Redmile-Gordon et al., 2013).

### Substrate Depletion method

Test compounds (substrates) were dissolved to make 10 mM stocks in DMSO. The microsome incubation mixtures (at different microsomal protein concentration 0.1, 0.25, 0.5, 0.75 and 1 mg/mL) was pre-incubated at 37 °C for 10 min, then placed on a Biomek liquid handling platform (Biomek i7 automated workstation, Beckman Coulter Life Sciences). Working substrate concentrations 500, 250, 100, 50, 10, 5, 1, 0.5 µM were prepared and added to 96-well plates containing the microsome incubation mixture (which consists of co-factors NADPH/UDPGA, microsomes, potassium phosphate buffer/tris-HCL phosphate buffer, MgCl, alamethicin) to start the reaction, with a final substrate concentration at 5, 2.5, 1, 0.5, 0.1, 0.05, 0.01, 0.005 µM. Aliquots were removed from the incubation reaction at 0, 5, 15, 30 and 45 min, timepoints and quenched with ice-cold acetonitrile containing labetalol internal standard. Assay samples were centrifuged for 15 min at 4000 rpm and supernatants were diluted 1:1 with water and analyzed by UPLC/MS/MS.

### Ultra-performance liquid chromatography (UPLC)

A Waters UPLC system was used to analyze the processed assay samples on an Acquity UPLC HSS T3 C18 column (2.1 x 50 mm, 1.8µm) which was maintained at 40 °C in a column oven. Mobile phase A consisted of water with 0.1 % formic acid, and mobile phase B consisted of acetonitrile with 0.1 % formic acid. The LC gradient is shown on Table 1.

**Table 1.** UPLC gradient conditions.

| **Time** | **Flow**<br>**(mL/min)** | **% A** | **% B** |
| --- | --- | --- | --- |
| Initial | 1.000 | 95.0 | 5.0 |
| 0.03 | 1.000 | 95.0 | 5.0 |
| 0.80 | 1.000 | 5.0 | 95.0 |
| 0.85 | 1.000 | 5.0 | 95.0 |
| 1.00 | 1.000 | 95.0 | 5.0 |
| 1.10 | 1.000 | 95.0 | 5.0 |

### Mass spectrometry (MS)

A triple quadrupole mass spectrometer (Waters Xevo TQS-micro) was used for detection and quantitation in an electrospray (ESI) positive and negative ion mode for all the compounds, internal standard, and the monitored metabolites. The capillary and cone voltage were 0.60 kV and 20 V respectively. The other parameters such as the desolvation and cone gas flow, the collision energy was 900 L/Hr, 20 L/Hr and 20 V respectively. The desolvation temperature applied was 650 °C. The test compounds were detected using multiple reaction monitoring, with compound specific MS methods (see Table 2).

**Table 2.**
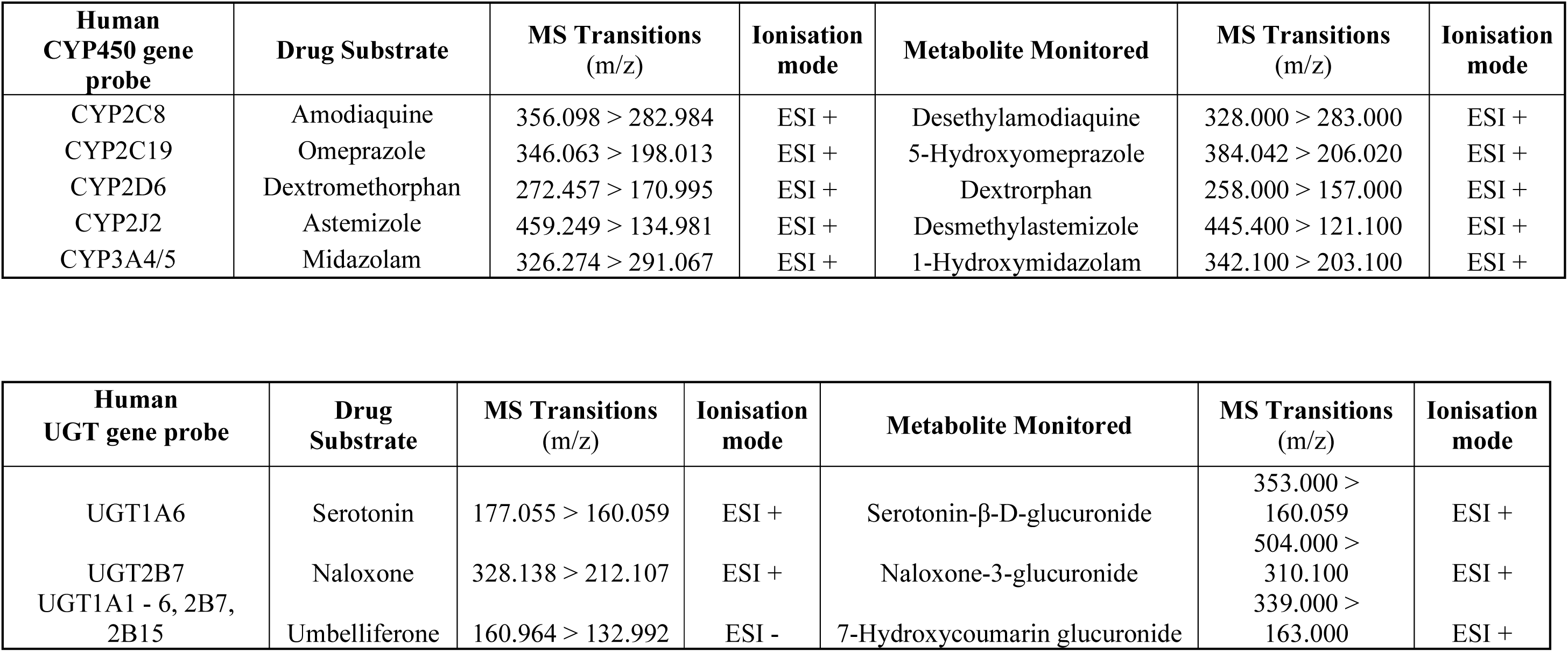
MS methods of CYP450 and UGT drug substrates and their corresponding marker metabolites.

| **Human CYP450 gene probe** | **Drug Substrate** | **MS Transitions (m/z)** | **Ionisation mode** | **Metabolite Monitored** | **MS Transitions (m/z)** | **Ionisation mode** |
| --- | --- | --- | --- | --- | --- | --- |
| CYP2C8 | Amodiaquine | 356.098 > 282.984 | ESI + | Desethylamodiaquine | 328.000 > 283.000 | ESI + |
| CYP2C19 | Omeprazole | 346.063 > 198.013 | ESI + | 5-Hydroxyomeprazole | 384.042 > 206.020 | ESI + |
| CYP2D6 | Dextromethorphan | 272.457 > 170.995 | ESI + | Dextrorphan | 258.000 > 157.000 | ESI + |
| CYP2J2 | Astemizole | 459.249 > 134.981 | ESI + | Desmethylastemizole | 445.400 > 121.100 | ESI + |
| CYP3A4/5 | Midazolam | 326.274 > 291.067 | ESI + | 1-Hydroxymidazolam | 342.100 > 203.100 | ESI + |

| **Human UGT gene probe** | **Drug Substrate** | **MS Transitions (m/z)** | **Ionisation mode** | **Metabolite Monitored** | **MS Transitions (m/z)** | **Ionisation mode** |
| --- | --- | --- | --- | --- | --- | --- |
| UGT1A6 | Serotonin | 177.055 > 160.059 | ESI + | Serotonin-β-D-glucuronide | 353.000 > 160.059 | ESI + |
| UGT2B7 | Naloxone | 328.138 > 212.107 | ESI + | Naloxone-3-glucuronide | 504.000 > 310.100 | ESI + |
| UGT1A1 - 6, 2B7, 2B15 | Umbelliferone | 160.964 > 132.992 | ESI - | 7-Hydroxycoumarin glucuronide | 339.000 > 163.000 | ESI + |

### Statistical analysis

Statistical analysis was performed using SPSS Statistics software (version 29.0.1.0, IBM). Compared means and proportions, independent-samples t test, and the computation of 95 % confidence intervals (Cls) were performed (Kapungu et al., 2020). The null hypothesis assumed no significant difference between the test (pig spleen microsomes) and control (pig liver microsomes) groups (Kapungu et al., 2020). One-Way ANOVA (analysis of variance) was performed on the initial rate of clearance observed in pig spleen and that observed in pig liver across the range of microsomal protein concentration ([MP]) of substrates omeprazole, astemizole, serotonin and amodiaquine datasets. The null hypothesis was rejected for all (p value > 0.05) with exception to the latter which had a p value < 0.05 indicating a difference between pig spleen microsomes and pig liver microsomes initial rate of clearance datasets across the different [MP].

## Results

### RNA-seq transcriptome analysis of CYP450 and UGT enzymes in Pig spleen and liver

Gene expression of CYP450 (Figure 2A) and UGT (Figure 2B) enzymes are confirmed in pig spleen by the NGS analysis. The metabolic genes identified showed differences in the expression levels between pig spleen to liver compared to human spleen. The CYP450 expression level is over 2-fold greater in pig liver to the spleen, but comparable expression levels are observed between the human spleen and pig spleen. Where CYP450 enzymes are the most predominant phase I enzymes in pig liver, aldehyde dehydrogenase seems to be the most expressed in both the pig and human spleen. However, significant differences were observed with the phase II enzymes particularly for the UGT enzymes. In pig liver, UGT is the second most expressed at 18 % compared to 0.1 % observed in pig spleen and 4 % in human spleen (see Figure 2B). Only two UGT genes were identified in pig liver and spleen (see Table 3); though the expression levels of phase II enzymes observed in pig spleen appear similarly to the human spleen.

**Figure 2:**
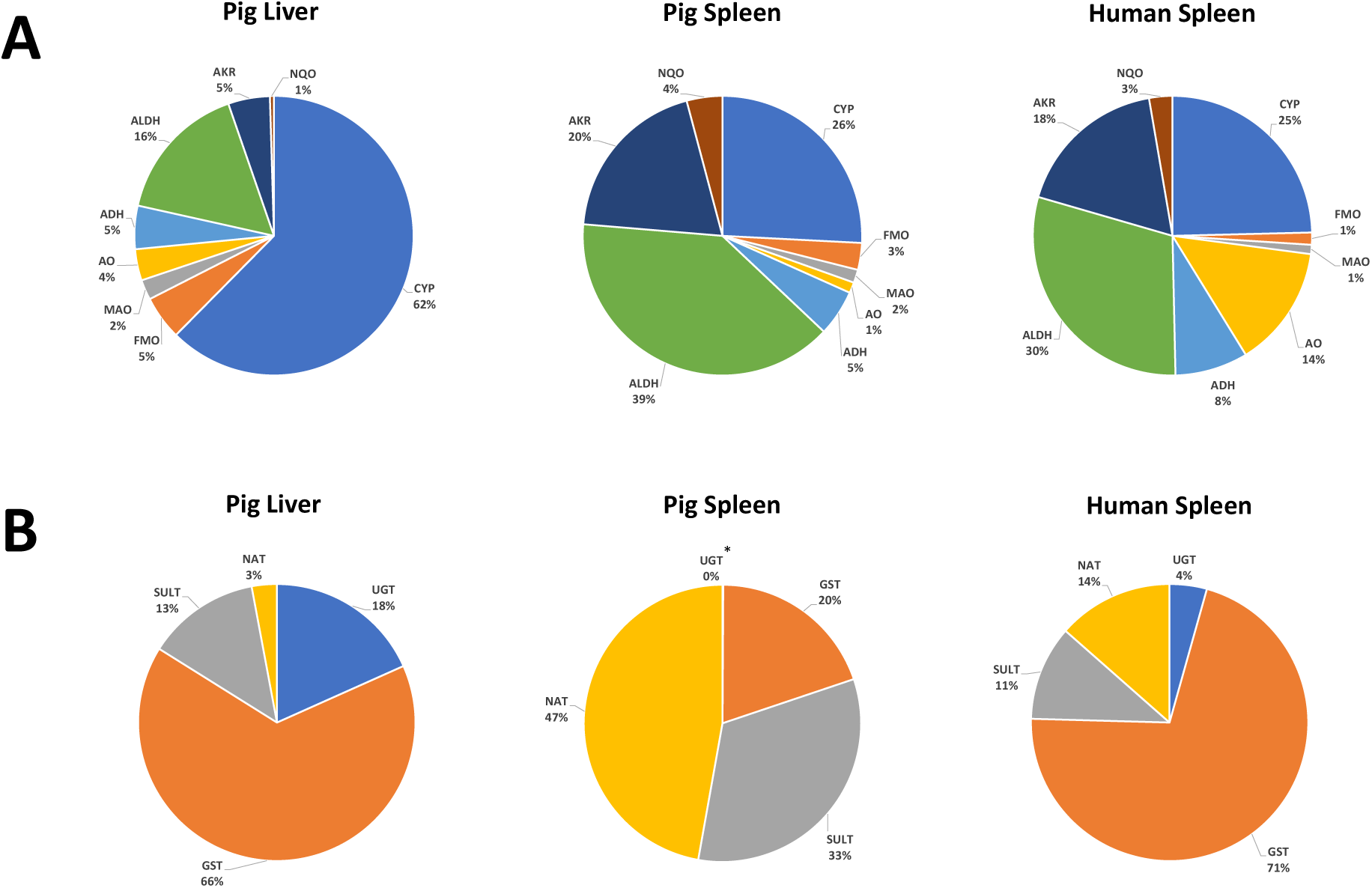
Gene expression of CYP450 (**A**) and UGT (**B**) enzymes in pig liver and spleen. Human spleen was used as reference sourced from ARCHS^4^ database. Abbreviations: Alcohol Dehydrogenases (ADH), Aldehyde Dehydrogenases (ALDH), Aldo-Keto Reductases (AKR), Aldehyde Oxidases (AO), Cytochrome P450 (CYP), Flavin-Containing Monooxygenases (FMO), Monoamine Oxidases (MAO), NADPH: Quinone Reductases (NQO), Glutathione S-Transferases (GST), Sulfotransferases (SULT), N-Acetyltransferases (NAT), Uridine Diphosphate Glucuronosyltransferases (UGT). *Pig Spleen UGT 0.1 %.

**Table 3.** The top 20 expressed CYP450 and UGT genes identified in pig by NGS analysis.

| Reference<br>CYP450 genes<br>in<br>Human Spleen | CYP450 genes in Pig |  | Reference UGT<br>genes in<br>Human Spleen | UGT genes in Pig |  |
| --- | --- | --- | --- | --- | --- |
|  | Pig Liver | Pig Spleen |  | Pig Liver | Pig Spleen |
| CYP20A1 | CYP3A29 | CYP1B1 | UGT3A2 | UGT2B31 | UGT8 |
| CYP51A1 | CYP2E1 | CYP27A1 | UGT8 | UGT8 | UGT2B31 |
| CYP1B1 | CYP2A19 | CYP2S1 | UGT1A1 |  |  |
| CYP4V2 | CYP2D25 | CYP20A1 | UGT2B17 |  |  |
| CYP2R1 | CYP27A1 | CYP51A1 | UGT2B4 |  |  |
| CYP3A5 | CYP1A2 | CYP2R1 | UGT1A6 |  |  |
| CYP2U1 | CYP2C42 | CYP2E1 | UGT1A7 |  |  |
| CYP2S1 | CYP4A24 | CYP4V2 | UGT2B11 |  |  |
| CYP7B1 | CYP51A1 | CYP7B1 | UGT3A1 |  |  |
| CYP46A1 | CYP4V2 | CYP2U1 | UGT1A3 |  |  |
| CYP27A1 | CYP7A1 | CYP2A19 | UGT2A1 |  |  |
| CYP1B1-AS1 | CYP7B1 | CYP3A29 | UGT2B10 |  |  |
| CYP2D7 | CYP1A1 | CYP2J34 | UGT2B15 |  |  |
| CYP4A22-AS1 | CYP2B22 | CYP4A24 | UGT1A10 |  |  |
| CYP4F12 | CYP4F8 | CYP39A1 | UGT2B7 |  |  |
| CYP4F3 | CYP2U1 | CYP2D25 | UGT2B28 |  |  |
| CYP2E1 | CYP39A1 | CYP11A1 | UGT2B27P |  |  |
| CYP2D6 | CYP2J34 | CYP26B1 | UGT2B25P |  |  |
| CYP1A2 | CYP8B1 | CYP2B22 | UGT1A5 |  |  |
| CYP4F8 | CYP26A1 | CYP46A1 | UGT2B24P |  |  |

Some of the major contributing CYP450 enzymes (or its equivalents) to drug metabolism in human liver are observed among the top expressed in both the human and pig spleen, and these include *CYP3A5/Cyp3a29*, *Cyp2e1*, *CYP2D6/Cyp2d25*, *CYP1A2*, *Cyp2a19* and *Cyp2j34*. It may appear that the pig spleen consists of more drug metabolizing CYP450s compared to that in the human spleen but in terms of relative abundance, from *Cyp2e1* down in the chronological order shown in Table 3, the genes are lowly expressed in pig spleen. Surprisingly, the two UGT genes identified in pig liver are not determined for their metabolic function though UGT1A, UGT2A and UGT2B are known to metabolize xenobiotics and endogenous compounds in humans (Bhatt et al., 2018). However, there is a difference of UGT expression levels between the pig liver (99.9 % UGT2B31, 0.1% UGT8) and spleen (negligible UGT2B31, almost 100% UGT8).

### Validation of CYP450 and UGT enzymes protein expression in Pig spleen and liver using western blot and immunofluorescence

Protein lysates isolated from fresh pig liver and spleen tissues were obtained from four individual animals and used for western immunoblotting. In addition, pig liver and spleen microsomes from the same animals were pooled and prepared via ultracentrifugation and semi-quantified to determine protein expressions. Particular attention was given to CYP450 and UGT genes identified by NGS. Strong bands indicating positive expression of the probed proteins (e.g. CYP450 generic and *Ugt8*) at the expected protein sizes of approx. 55 kDa, were observed in all the pig liver lysate samples (See Figure 3A). Faint bands expressing CYP450 generic were only observed in pig spleen microsomes sample and two pig spleen lysate samples. In contrast, low to moderate intensity bands expressing *Ugt8* were observed in all four spleen lysate samples, but not in the spleen microsomes sample. We have highlighted for both CYP450 generic and *Ugt8* in Figure 3A using black arrows to emphasis where the original faint bands are located.

**Figure 3:**
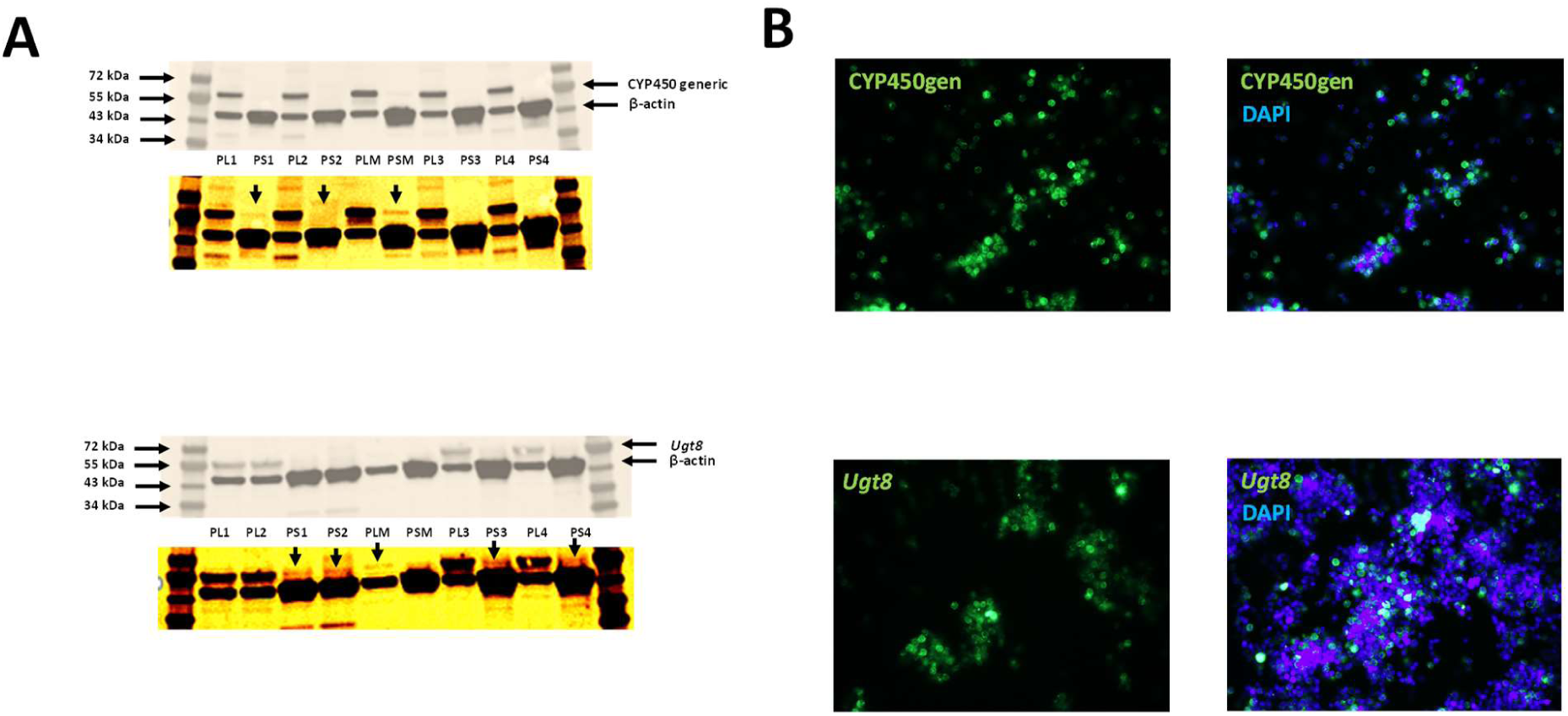
Expression levels of CYP450 and UGT enzymes in **A.** Western Blot analysis of pig liver (PL1-4) and pig spleen (PS1-4) protein lysates and microsomes with β-actin as the loading control **B.** Immunofluorescence Alexa Fluor 488 stain using pig splenocytes. PML: pooled microsomes from liver tissue; PMS: pooled microsomes from spleen tissue.

An indirect immunofluorescence labelling was carried out in pig hepatocytes and splenocytes to confirm the protein expression observed from the western blot analysis. The immunofluorescence images in Figure 3B showed CYP450 generic and *Ugt8* protein expression. Based on the intensity and abundance of the fluorescent signal, a moderate protein expression of CYP450, and a low-to moderate expression of *Ugt8* was observed. The use of a non-antibody stain (Dapi) was used as an additional control to indicate where the protein expression is localized.

### Microsomal drug disposition (clearance and metabolite formation)

Selected CYP450s and UGTs representing the major contributing drug metabolizing enzymes known in human liver were investigated to determine the drug elimination rate in pig liver and spleen. The substrate depletion approach was adopted in an assay to monitor the disappearance of drug substrates over time as probes for the target CYP450 and UGT enzymes (See Table 2). The resulting formation of metabolites was also monitored as confirmation that drug metabolism in pig spleen has occurred. The initial rate profiles in pmoles/min/mg of protein depicted in Figure 4 were normalized intrinsic clearance (Clint) with mg of protein concentration (See Supplementary data Table 1.0 - 4.1).

**Figure 4:**
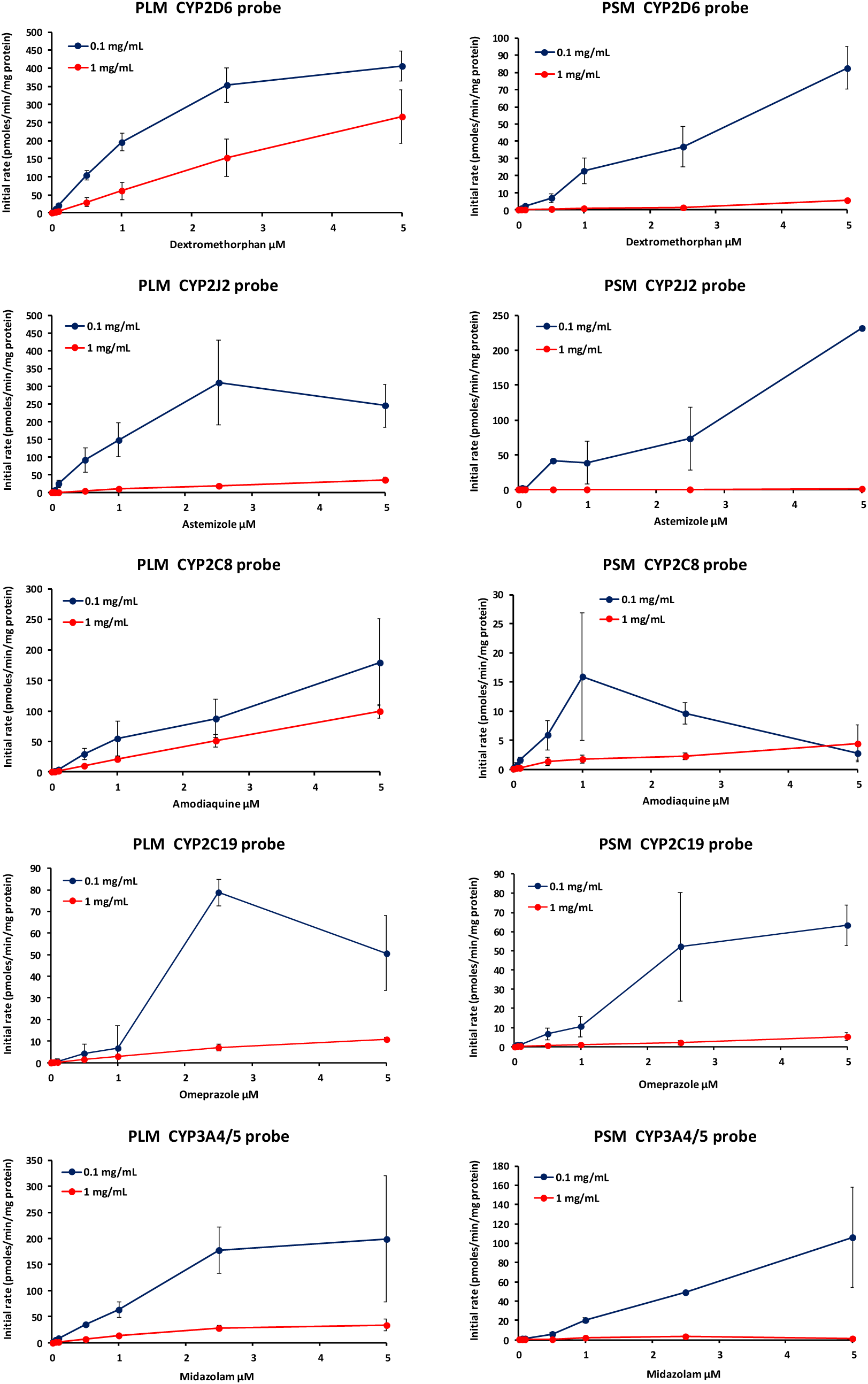
CYP450 graph profiles at 0.1 mg/mL and 1 mg/mL protein concentration in pig liver and spleen microsomes determining the initial rate of clearance of five drug substrates (Dextromethorphan, Astemizole, Amodiaquine, Omeprazole and Midazolam). PLM (pig liver microsomes) and PSM (pig spleen microsomes)

Five CYP-mediated metabolites were detected in spleen microsomes suggesting positive confirmation of drug metabolism and apparent CYP450 activity in pig spleen. All but one drug substrate (astemizole) in pig spleen microsomes showed an increase in drug clearance proportional to an increasing drug substrate concentration. With the substrate, astemizole, a sharp decrease in drug elimination was observed from 2.5 µM. The drug clearance overall was lower in pig spleen microsomes compared to the drug clearance observed in pig liver microsomes. However similarly in pig liver microsomes, it appears that the elimination of drugs is predominant at lower microsomal protein concentration as in pig spleen microsomes (See Figure 4).

The peak area response indicating the CYP450 metabolites’ levels were obtained from the mass spectrometry analysis monitoring for the specific metabolites. In addition, the metabolite response profile was displayed as a function of time and microsomal protein concentration (See Figure 5). Metabolites (Dextrorphan and 1-Hydroxymidazolam) were only detected from 1 µM at both 0.1 and 1 mg/mL microsomal protein concentration including at 5 and 45 minutes. This was also comparable to the metabolite profile of desmethylastemizole in 0.1 mg/mL microsomal protein concentration at both 5 and 45 minutes. The other metabolites (Desethylamodiaquine and 5- Hdroxyomeprazole) showed a similar profile compared to the same metabolite profile observed in pig liver microsomes. At 1 mg/mL microsomal protein concentration, higher metabolite levels of desmethylastemizole were observed compared to 0.1 mg/mL, though appeared to stabilize by 2.5 µM substrate concentration of astemizole. In general, metabolite response levels in pig spleen microsomes were comparable between 5 and 45 minutes, with the latter timepoint just ahead in metabolite levels. The variation across timepoints is significant for some of the observed results, however the apparent drug metabolism observed in pig liver microsomes (as the control tissue) for the tested drug substrates indicate a robust assay was conducted.

**Figure 5:**
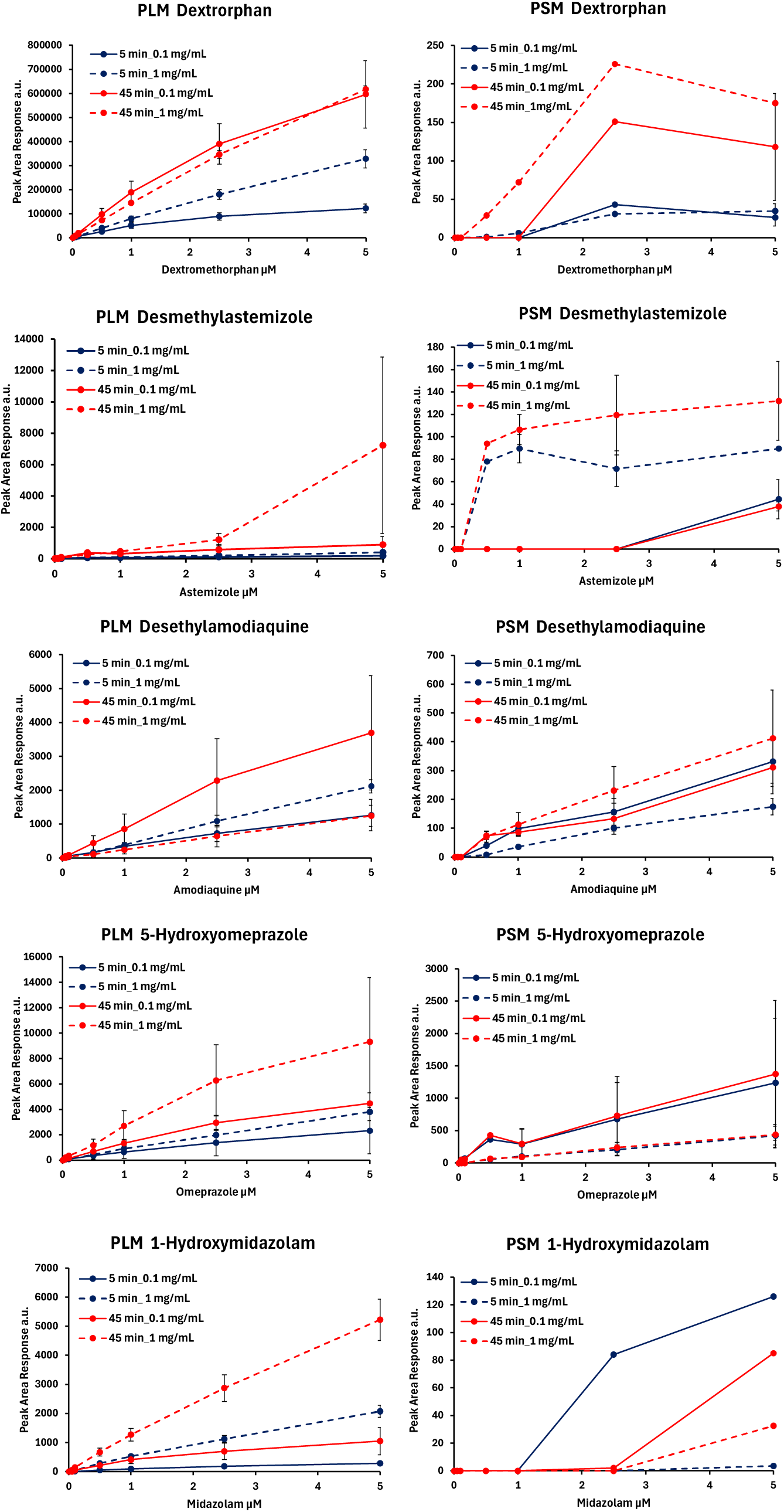
CYP450 graph profiles at 0.1 mg/mL and 1 mg/mL protein concentration in pig liver and spleen microsomes of the corresponding monitored metabolites formed.

The UGT-mediated rate of drug clearance also appeared to increase with increasing drug substrate concentration, with exception at 5µM for the substrate, umbelliferone. Somewhat similar with CYP450 profiles describing the rate of drug elimination, there is a clear difference between 0.1 and 1 mg/mL microsomal protein concentration in pig spleen microsomes (See Figure 6). The UGT-mediated drug clearance at 1 mg/mL protein concentration appears greater for the tested drug substrates than that observed generally for the CYP450-mediated drug clearance in pig spleen microsomes.

**Figure 6:**
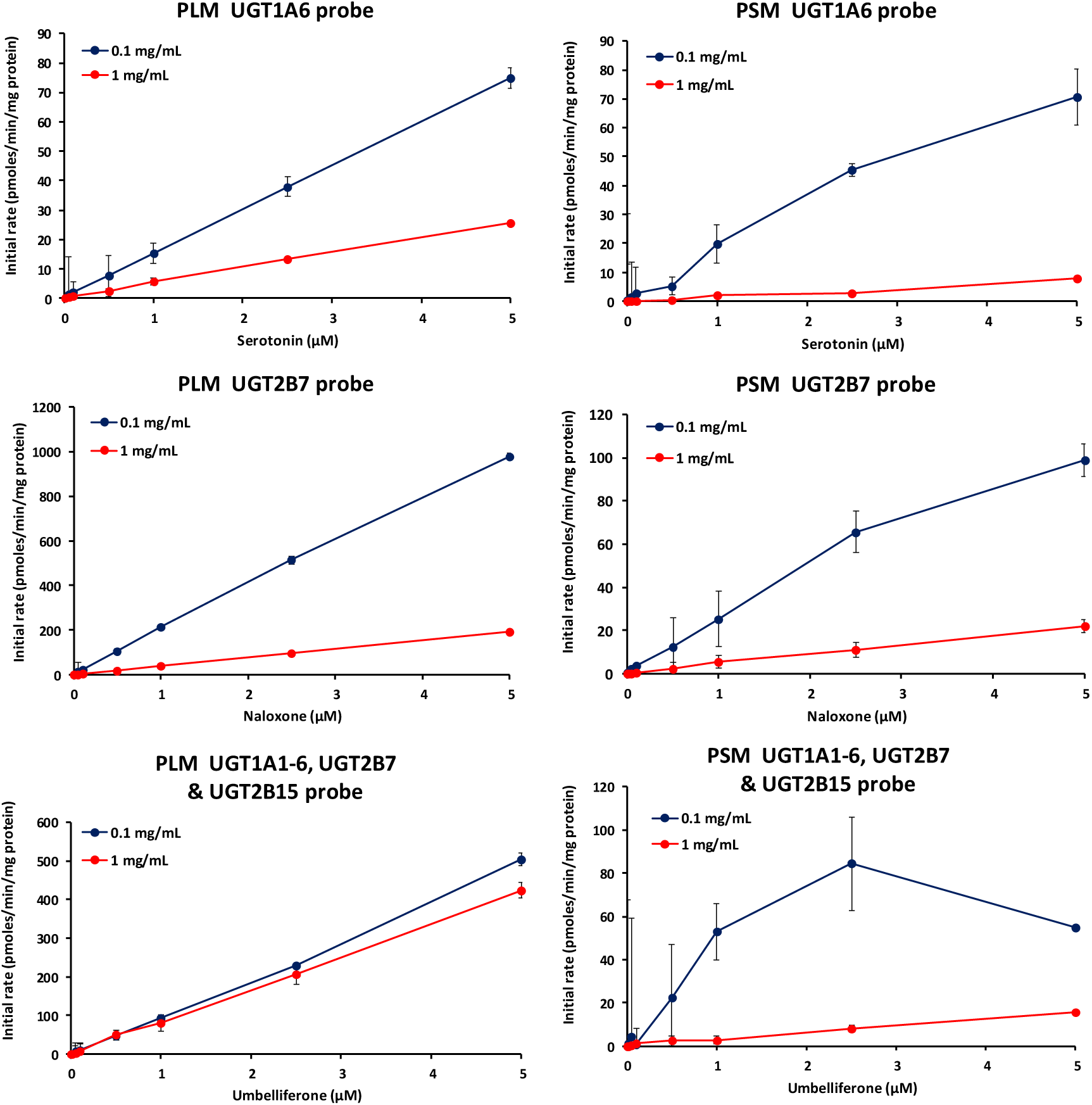
UGT graph profiles at 0.1 mg/mL and 1 mg/mL protein concentration in pig liver and spleen microsomes determining the initial rate of clearance of three drug substrates (Serotonin, Naloxone, and Umbelliferone). PLM (pig liver microsomes) and PSM (pig spleen microsomes)

The glucuronide metabolites were detected as low as ≤ 1 µM in both 0.1 and 1 mg/mL microsomal protein concentration and at 5 and 45 minutes. Positive glucuronidation reaction is indicated in pig spleen microsomes, with a reasonable level of formed metabolites detected from a small amount of drug substrate concentration (See Figure 7). Serotonin-β-D-glucuronide showed comparable levels of the metabolite at 5 and 45 minutes in 0.1 mg/mL microsomal protein concentration. Though, higher levels of the metabolite were observed at 45 minutes compared to 5 minutes in 1 mg/mL microsomal protein concentration. In contrast for Naloxone-3-glucuronide, there is a difference of metabolite levels between 5 and 45 minutes but significant at 45 minutes in 1 mg/mL microsomal protein concentration. Higher metabolite levels of 7-hydroxycoumarin-glucuronide were observed at lower microsomal protein concentration (0.1 mg/mL) compared to 1 mg/mL, but with comparable levels at both 5 and 45 minutes.

**Figure 7:**
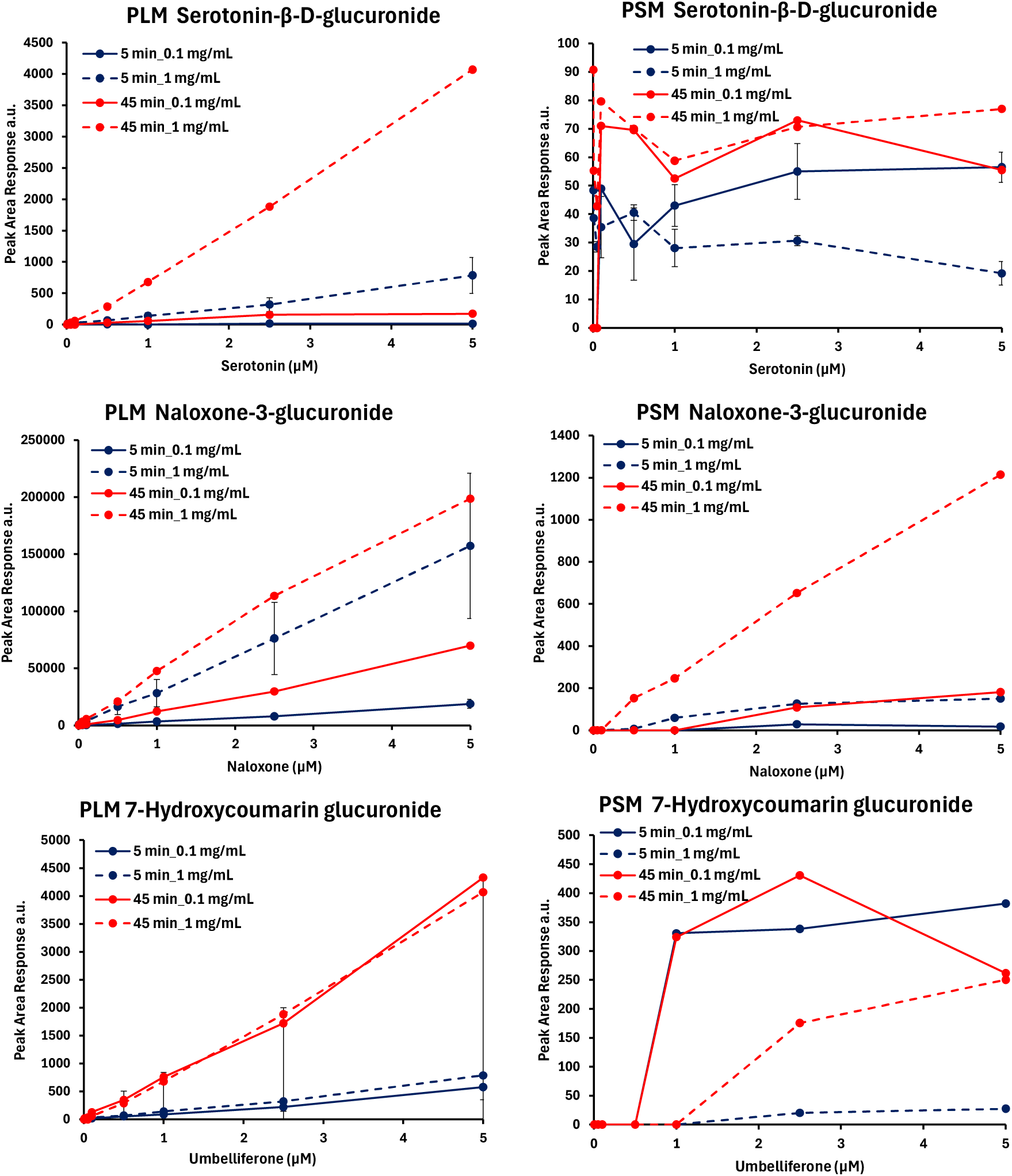
UGT graph profiles at 0.1 mg/mL and 1 mg/mL protein concentration in pig liver and spleen microsomes determining the initial rate of clearance of three drug substrates (Serotonin, Naloxone, and Umbelliferone). PLM (pig liver microsomes) and PSM (pig spleen microsomes)

As observed with the assay examining CYP450 activity, variation across the timepoints monitoring UGT activity was also apparent for some of the results, but the working conditions of the assay generally appeared not to be affected.

### Cellular drug disposition (clearance and metabolite formation)

The same drug substrates of CYP450 enzymes were used in a similar assay using pig hepatocytes and splenocytes. Cells contain both phase I and II enzymes and the substrate depletion cell-based assay was used to imitate the *in-vivo* condition and to determine the drug elimination rate in pig cells. The initial rate profiles in pmoles/min/0.5 million cells depicted in Figure 8 were normalized intrinsic clearance (Clint) with the number of cells in the incubation tested at 0.5 million cells (See Supplementary data Table 5.0 - 6.1).

**Figure 8:**
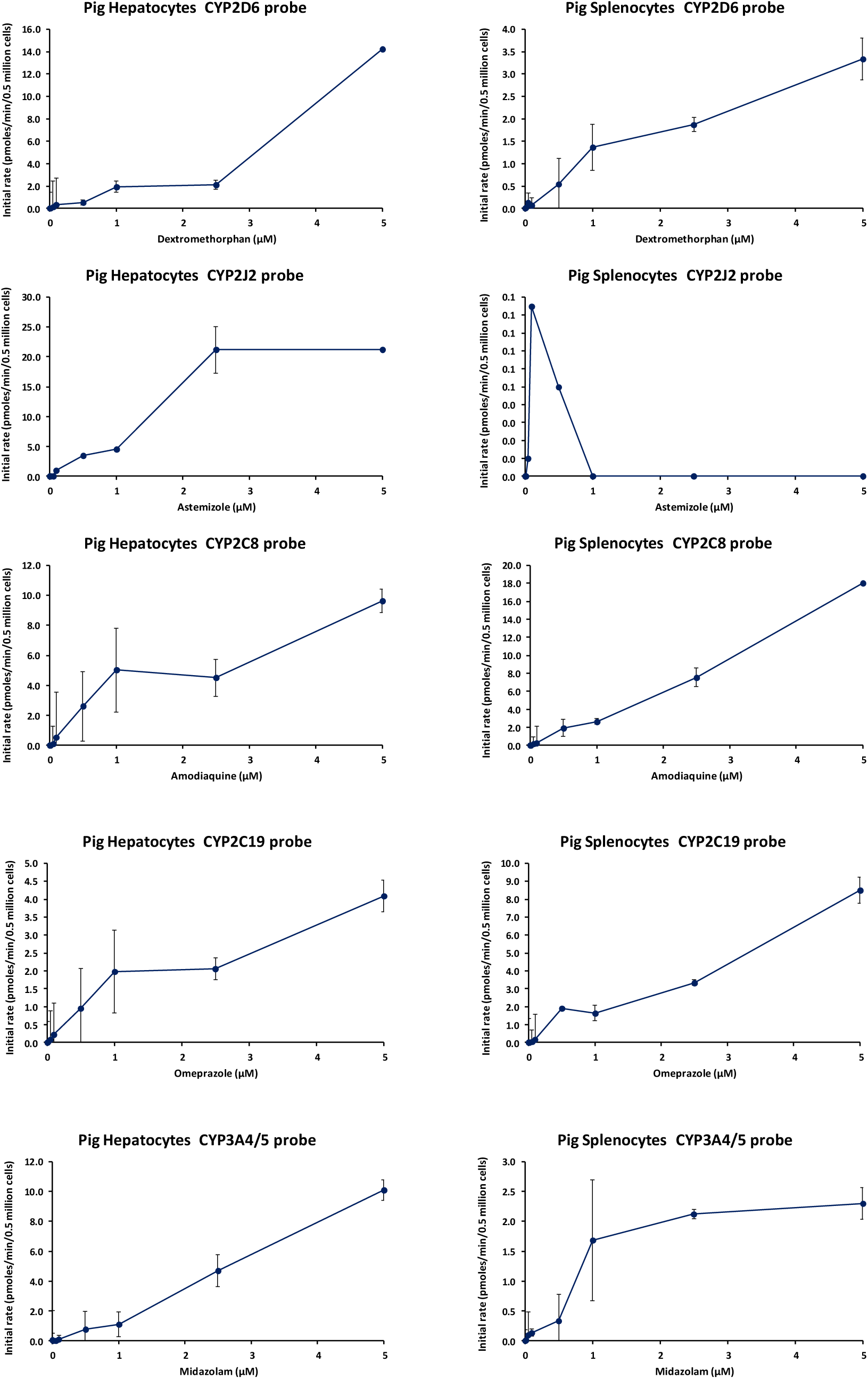
CYP450 graph profiles in pig hepatocytes and splenocytes determining the initial rate of clearance of five drug substrates (Dextromethorphan, Astemizole, Amodiaquine, Omeprazole and Midazolam), using a Michalis Menten model to display a linear regression curve.

**Figure 9:**
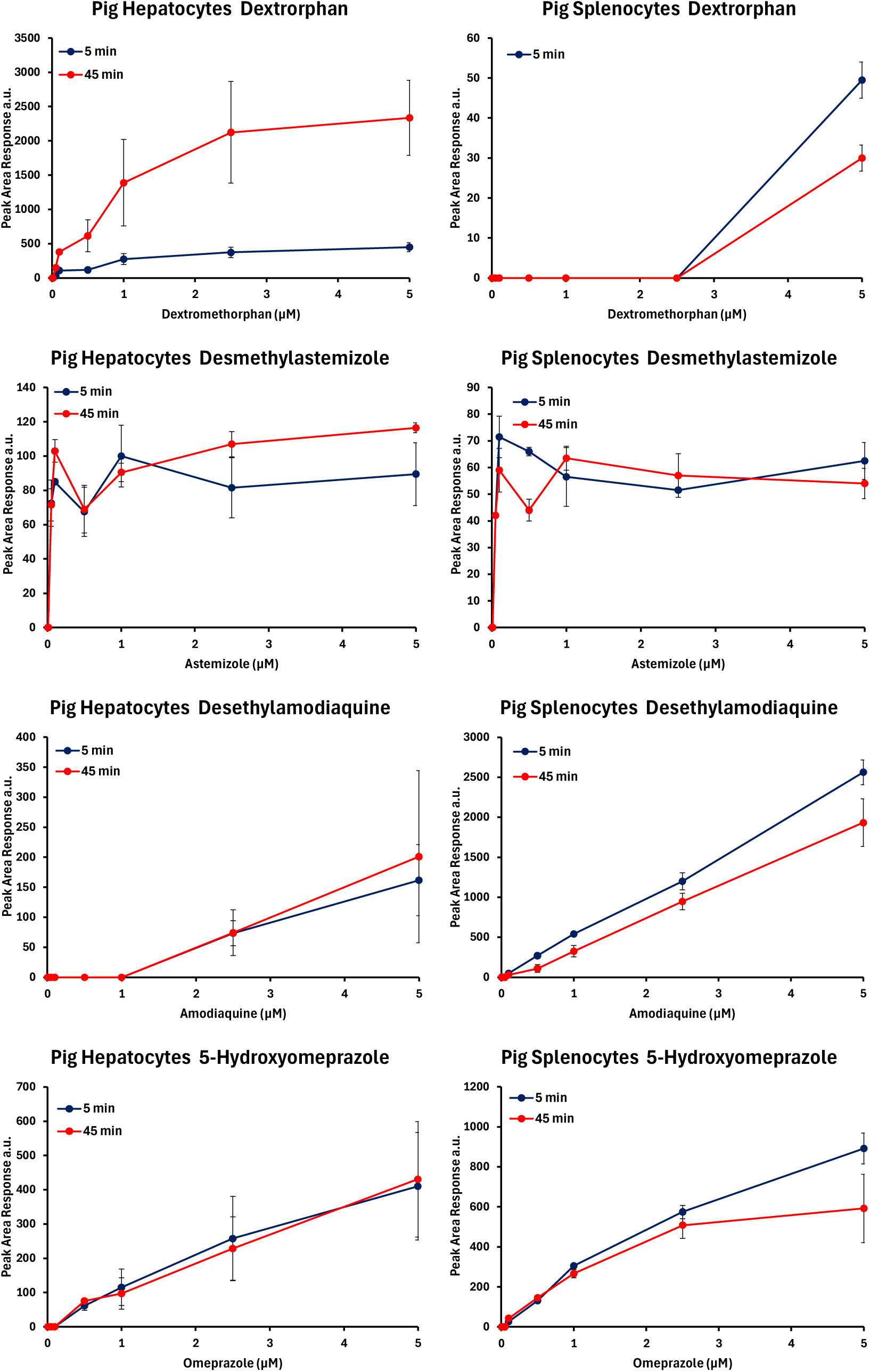
CYP450 graph profiles in pig hepatocytes and splenocytes illustrating the corresponding monitored metabolites formed.

CYP450-mediated monitored drug clearance appeared to increase proportionally to increasing drug substrate concentration with exception of the substrates, astemizole and midazolam. Drug elimination was observed up to 0.1 µM for astemizole and drug saturation from 2.5 µM was observed for midazolam. The rate of drug clearance overall appeared somewhat linear.

The metabolite formation was also monitored in pig hepatocytes and splenocytes to provide strong confirmation of drug metabolism observed in pig cells. Both metabolites in pig splenocytes, desethylamodiaquine and 5-hydroxyomeprazole showed similar profiles compared with the same metabolite profiles observed in pig hepatocytes. Moreover, significant metabolites levels of both were also observed in pig splenocytes compared to that observed in pig hepatocytes. Dextrorphan appeared to be only detected at 5 µM, whereas desmethylastemizole levels remain constant from 0.1 µM. No metabolite formation levels were detected for the drug substrate midazolam.

Likewise, both analysis monitoring drug clearance and metabolite formation in pig hepatocytes was successfully achieved and provided the acceptance criteria for the analysis in pig splenocytes.

## Discussion

The gene expression analysis revealed the genetic make-up of the spleen tissue and showed the abundance of mRNA sequences (key for making functional gene products) that included metabolic CYP450 and UGT genes. This analysis was performed to support the notion that the spleen should have CYP450 and UGT genes in its DNA to have the ability to metabolize drugs; since drugs are the main therapeutic tools that can be used to establish the therapeutic aims. Nonetheless, drug research studies have always involved using appropriate animal models for human, and the phylogenetic analysis based on mRNA sequences has shown close homology between pig to human. Such provided the foundation for selecting these models as appropriate for downstream analysis in this study, as having access to human tissues or even tissues from monkey are limited and have linked ethical implications. The equivalent CYP450 and UGT genes were subsequently identified from the RNA-seq transcriptome (via the NGS analysis) in untreated or basal pig spleens. Positive protein expression is a true representation of functional genes as they mainly carry out the biological processes in the body (Alberts et al., 2002). Confirmation of the protein expression of selected CYP450s and UGTs, as identified by NGS, in pig spleens would suggest two things, that those identified genes are not benign or non-coded genes and that the spleen tissue indeed contains functional CYP450 and UGT proteins.

What stands out generally from these functional observations in pig liver and spleen microsomes is that drug elimination appears to be higher at lower microsomal protein concentration (0.1 mg/mL) compared to a greater microsomal protein concentration at 1 mg/mL. It may seem logical to rationalize that increasing the protein concentration, increases the volume of available CYP450 or UGT enzymes for the metabolizing reactions. However, these observations appear contrary to that understanding and may likely be a result of two things: firstly, the reaction has reached saturation given that CYP450 enzymes have multiple binding sites (to account for its diverse substrate specificities capability) and the drug substrate amount has been used up at 0.1 mg/mL microsomal protein concentration. Increasing the microsomal protein concentration will not cause any changes as there is limited substrate available to occupy all the binding or active sites of the enzyme and this can be demonstrated from the profiles curving by 5 µM drug substrate concentration at 0.1 mg/mL; secondly, since the Clint values were normalized with mg of microsomal protein, this may indicate that there was less drug metabolism that took place initially and this profile is a result of non-specific binding of the drug substrates to other proteins in the microsome incubation.

Although the initial rate across the drug substrates in the cell-based assay were significantly lower compared to that in microsomes, it is worth mentioning that the drug substrates are exposed to phase I and II enzymatic reactions. In addition, the low drug elimination observed indicates that CYP450-mediated metabolism is not the major route of drug metabolism for these substrates in the cellular matrix. However, the metabolite formation profiles appear significantly higher particularly from the drug substrates amodiaquine and omeprazole in pig splenocytes. This suggest that the CYP450 and possibly UGT enzymes in pig splenocytes are much more efficient in metabolizing drugs in the cellular matrix compared to splenic microsomes. Moreover, the metabolite 1-hydroxymidazolam was not detected in either pig hepatocytes and splenocytes but was detected in both liver and spleen microsomes. It is important therefore to understand that the metabolites formed may be possible substrates for other enzymes or subject to glucuronidation by UGT enzymes, depleting the available metabolite formed before it is measured (See Figure 10). This can be compared against other metabolites such as desethylamodiaquine, 5- hydroxyomeprazole and desmethylastemizole given their higher detectable levels as they are not known to be substrates for other enzymes (See Figure 11).

**Figure 10:**
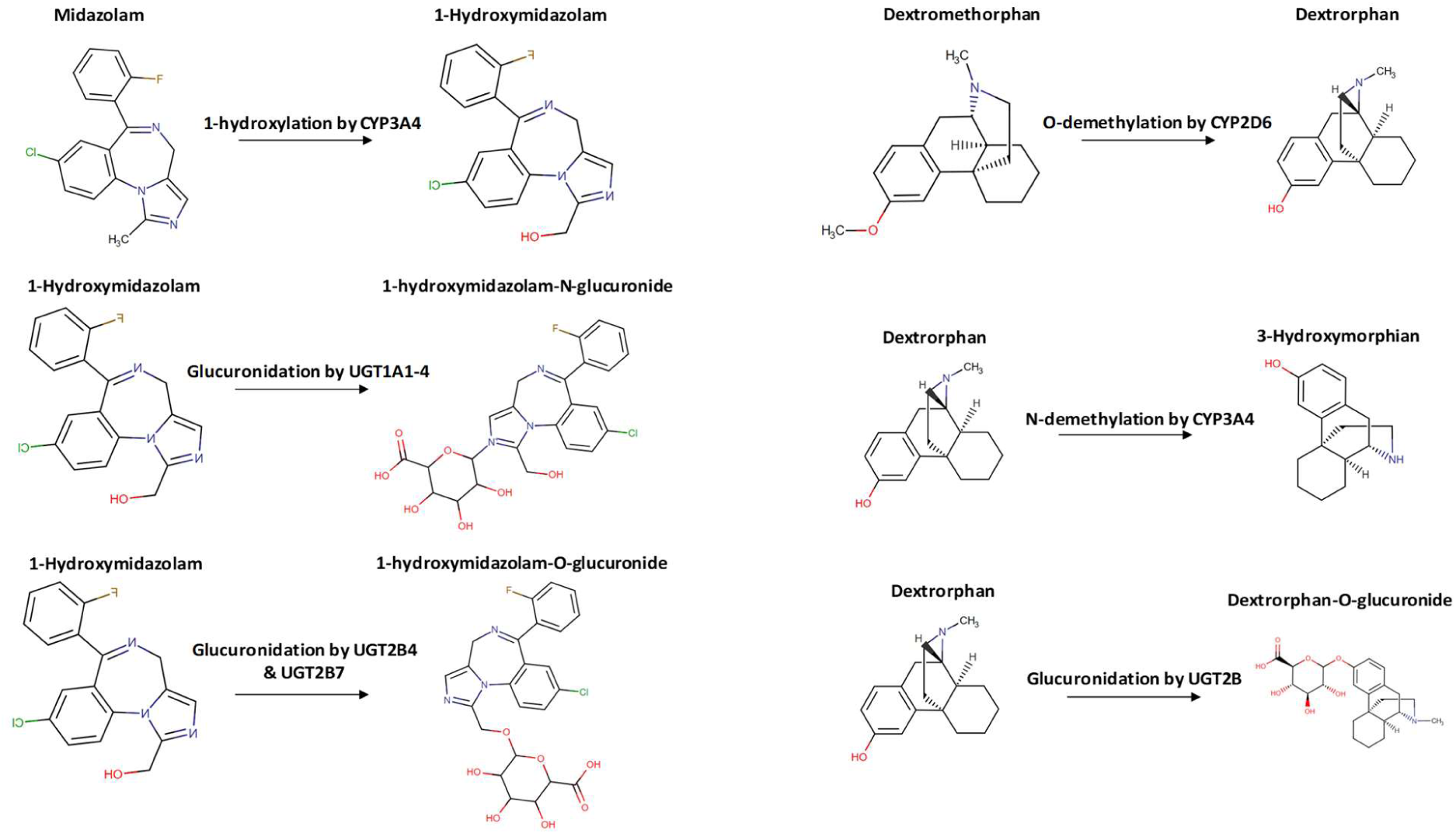
Examples of structures of CYP450 drug substrates, midazolam and dextromethorphan with their metabolites, 1-hydroxymidazolam and dextrorphan, including the type of metabolism

**Figure 11:**
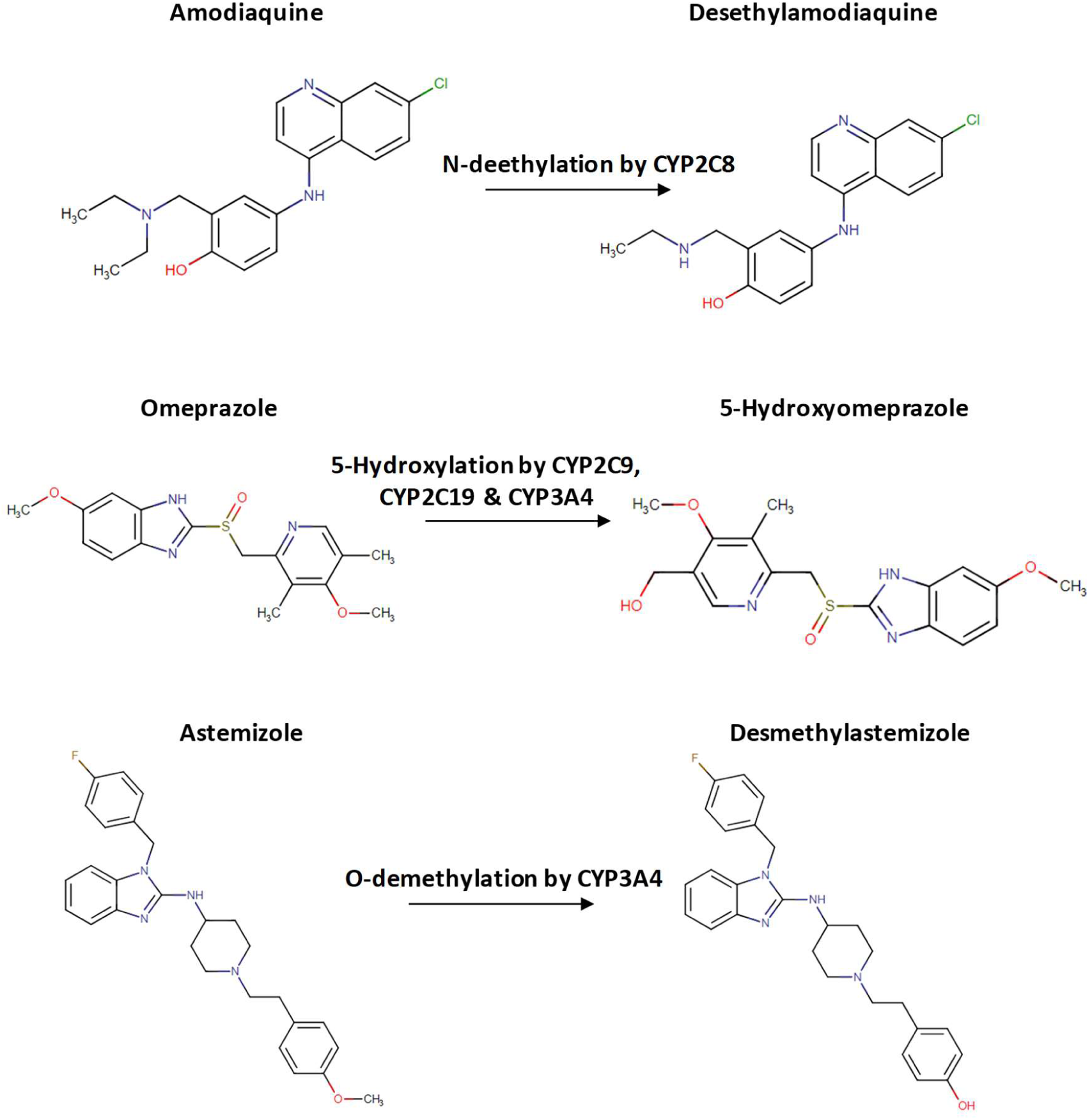
Examples of structures of CYP450 drug substrates, amodiaquine, omeprazole and astemizole with their metabolites, desethylamodiaquine, 5-hydroxyomeprazole and desmethylastemizole, including the type of metabolism.

Overall, the spleen appears to be an attractive immunotherapeutic target since administered drugs can access the immune cells area (periarteriolar lymphoid sheath) without facing challenging barriers based on its anatomy. The genetic, protein and functional expression of metabolic enzymes like the CYP450 and UGT enzymes have demonstrated that the spleen can process therapeutic drugs and perhaps much more efficient than the liver tissue.

In hindsight, the recommendations for future drug metabolism studies in the spleen would be to discover drug substrates specific to the CYP450 and UGT enzymes in the spleen. It would also be beneficial to subsequently perform an enzyme progress (saturation) curve to determine the Michalis Menten parameters (𝑉_max_ and 𝑘_m_) for those substrates in the spleen. In addition, it appears that a cell-based assay is a better option for determining enzyme kinetics in the spleen than using splenic microsomes and would eliminate the need to test for microsome drug binding in monitoring its impact on the enzyme’s kinetics. An *in-vivo* PK assay is necessary for splenic therapeutic intervention in a pre-clinical setting and should also be implemented to correlate with the data obtained from *in-vitro* studies.

To conclude, the spleen appears to be highly efficient in metabolizing drugs, requiring limited amount of drug substrate while detectable metabolite levels as low as 10 nM are observed. Overall, this study depicts the activities of CYP450s and UGTs in pig spleen to an extent and has demonstrated its potential for future therapeutic uses or manipulations.

### Author Contributions

CR, SP, PI conceived and designed research. PI performed experiments and analyzed data. CR, SP, PI interpreted results of experiments. CR, PI prepared figures. PI drafted manuscript. PI, CR edited and revised manuscript. PI, CR, SP approved the final version of manuscript.

## Funding

The work was funded by UKRI - BBSRC DTP iCase and Sygnature Discovery (grant RS3828 to SP, CR).

### Data availability

Data is provided as supplementary material.

