## Supplementary material for "The drug disposition in the Spleen: A novel study of CYP and UGT-mediated drug metabolism": Data

### SUPPLEMENTARY INFORMATION

**Table. 1.0	CYP450-mediated calculated Intrinsic clearance including ± standard deviation of 5 drug substrates in pig liver microsomes**

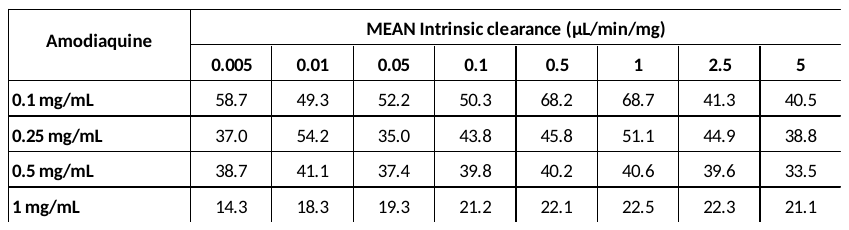

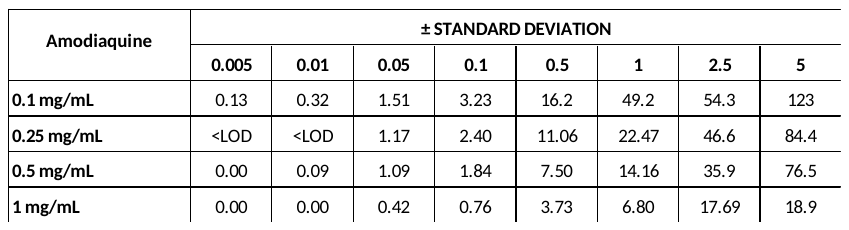

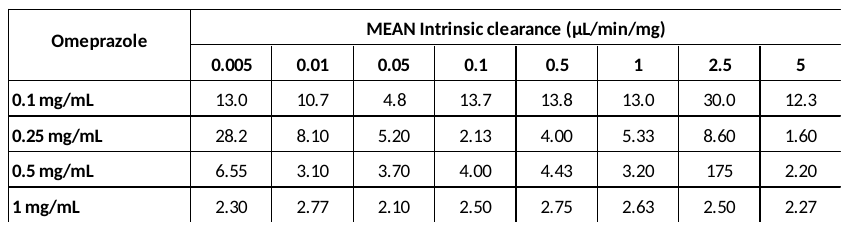

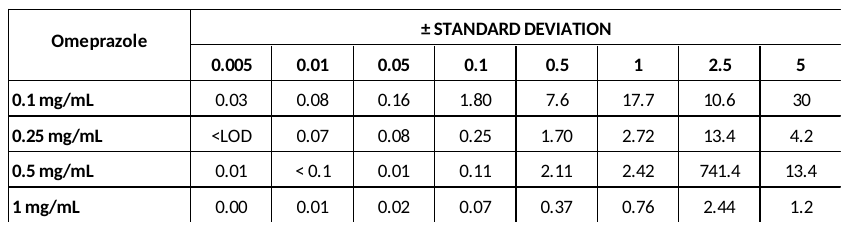

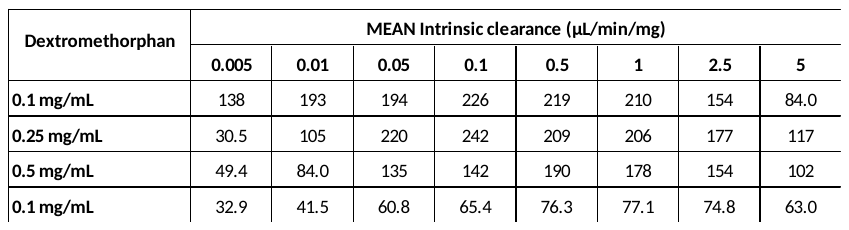

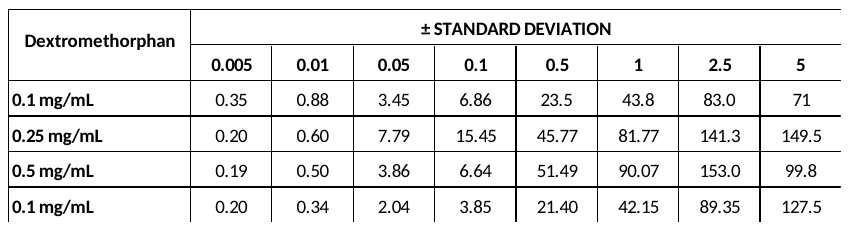

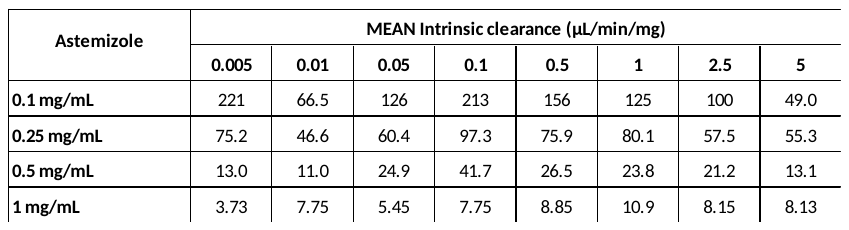

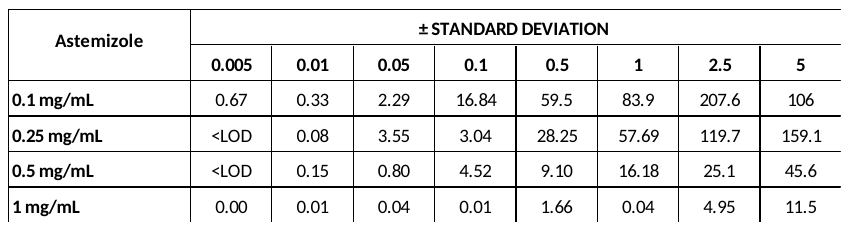

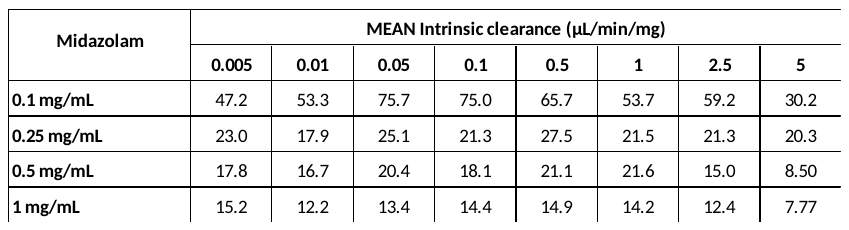

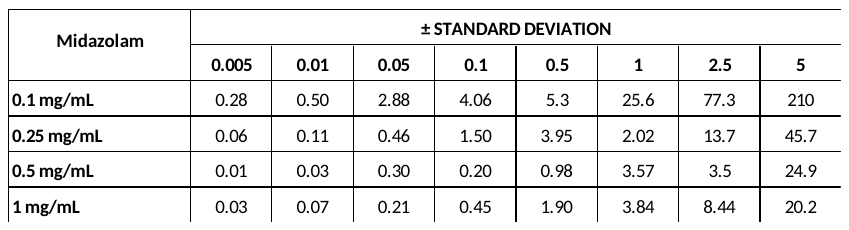

**Table. 1.1	MEAN CYP450-mediated drug elimination rate** $\boldsymbol{k}$ **(- gradient) and ± standard error for 5 drug substrates in pig liver microsomes**

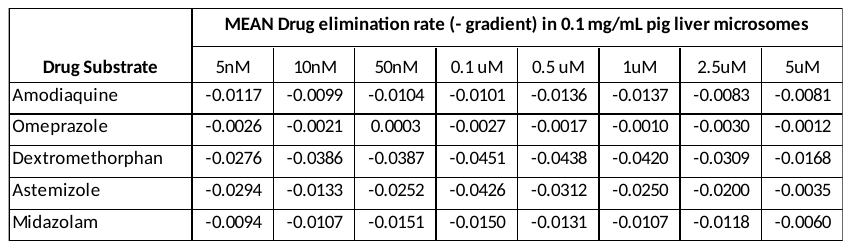

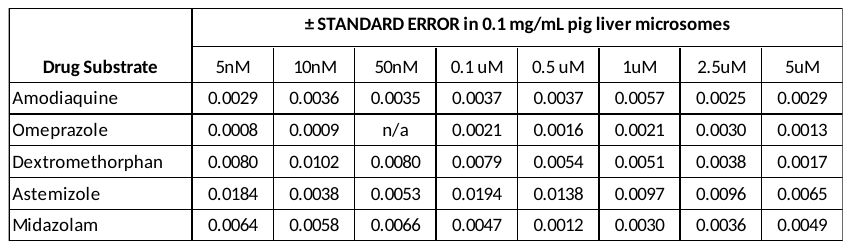

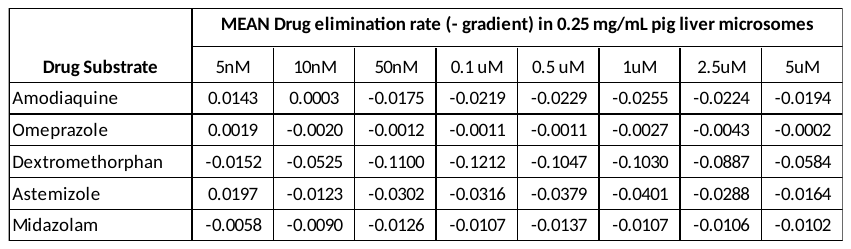

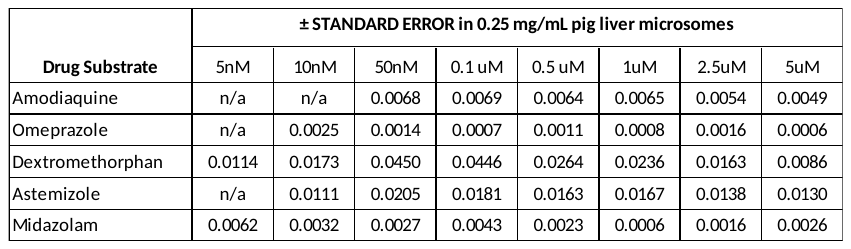

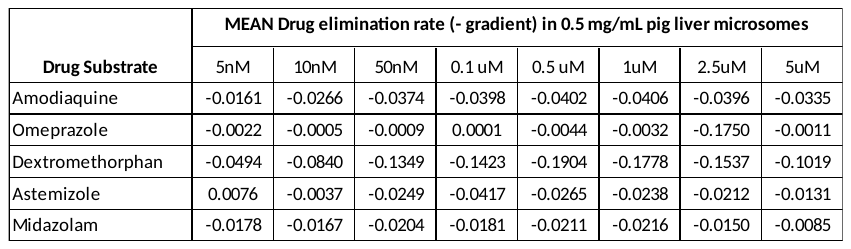

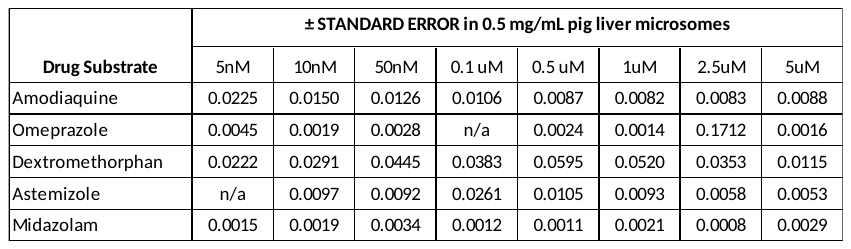

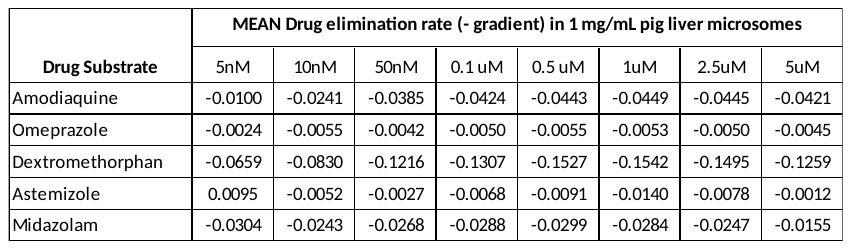

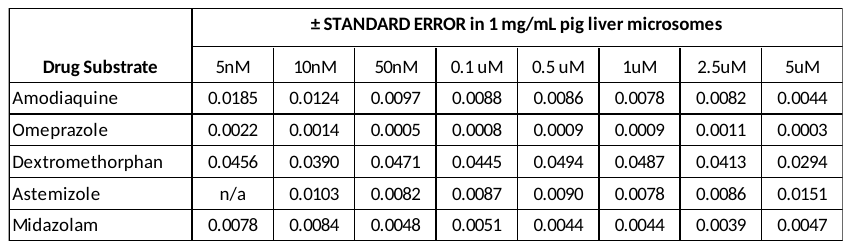

**Table. 2.0	CYP450-mediated calculated Intrinsic clearance including ± standard deviation of 5 drug substrates in pig spleen microsomes**

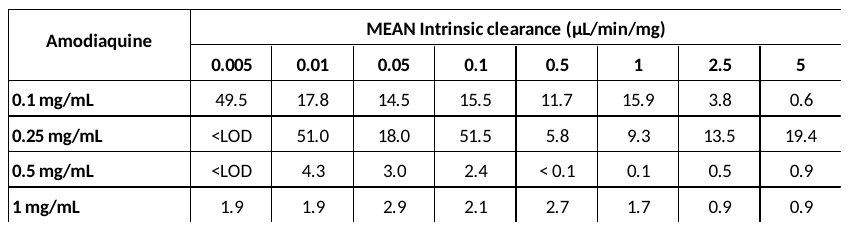

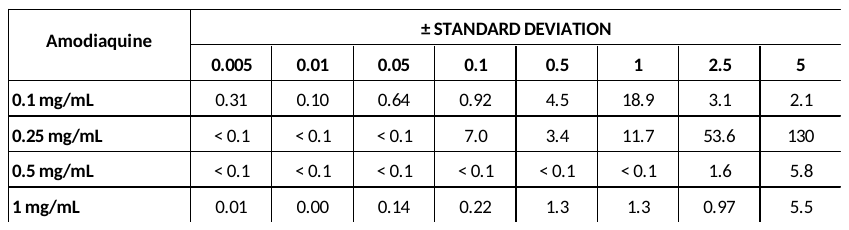

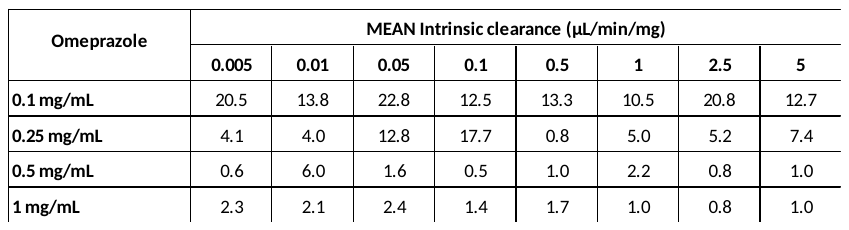

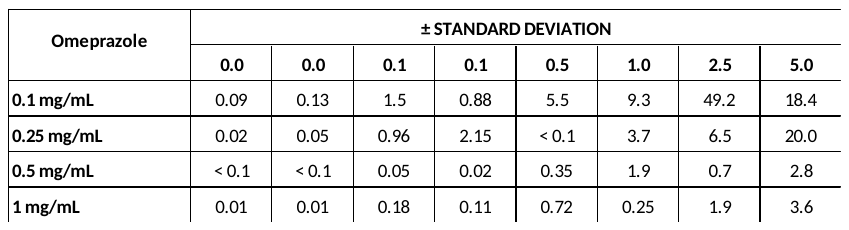

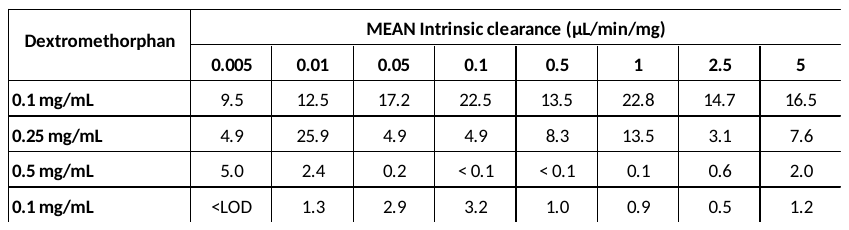

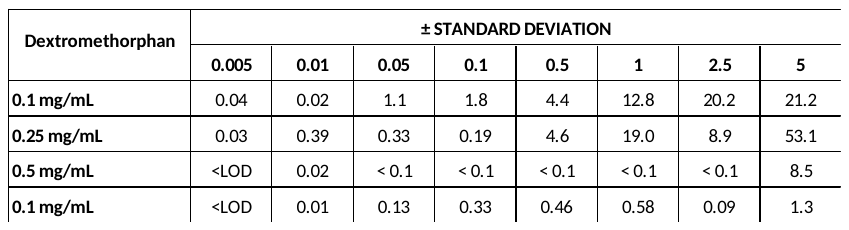

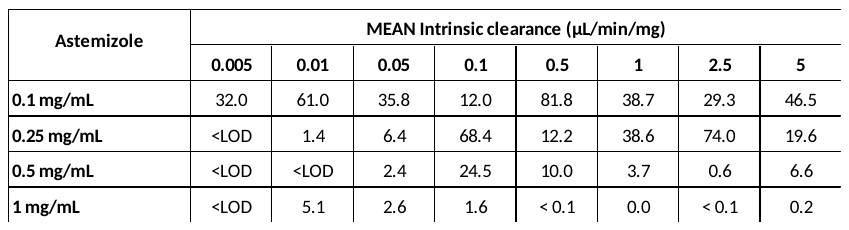

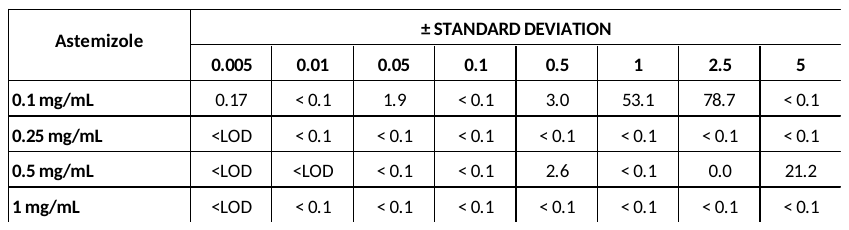

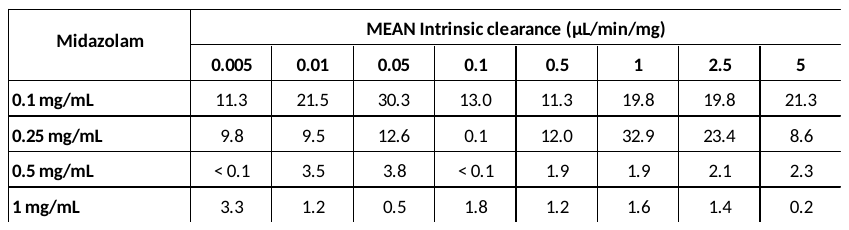

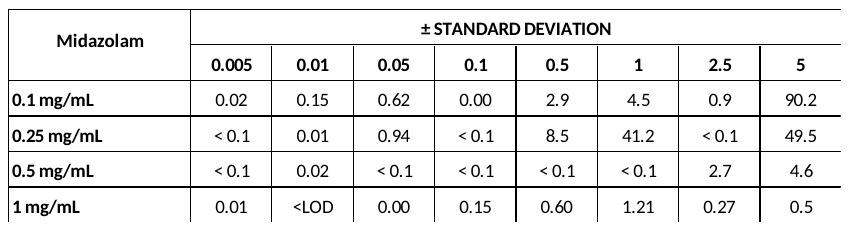

**Table. 2.1	MEAN CYP450-mediated drug elimination rate** $\boldsymbol{k}$ **(- gradient) and ± standard error for 5 drug substrates in pig spleen microsomes**

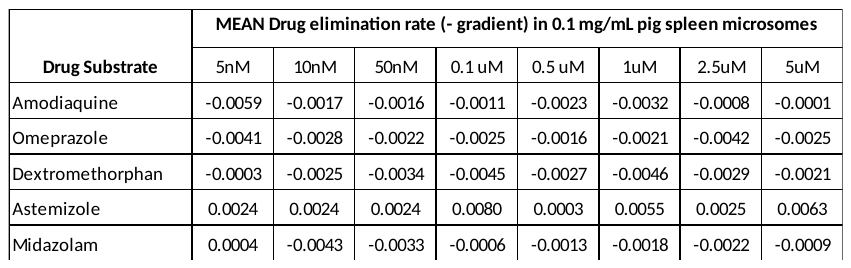

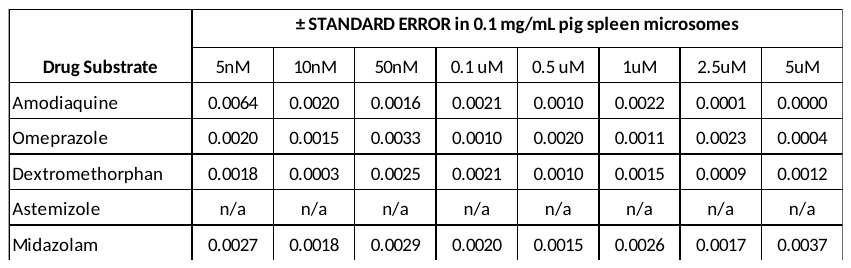

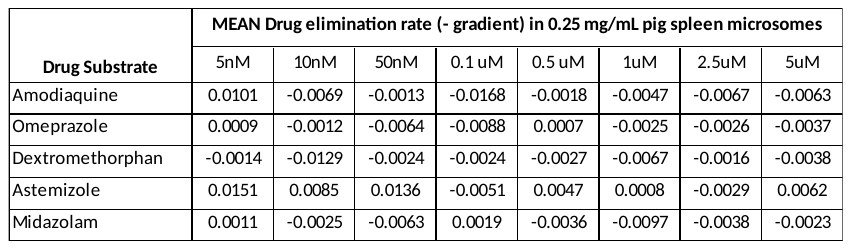

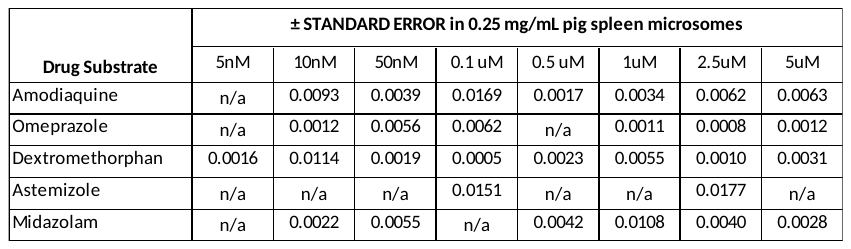

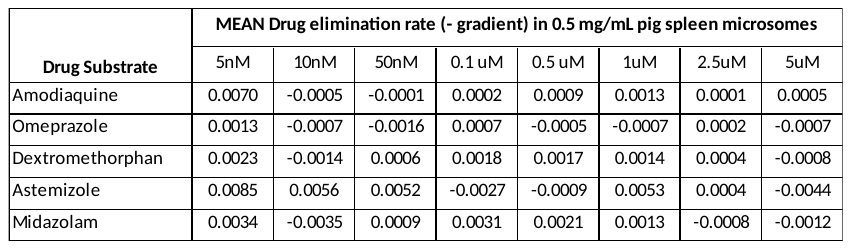

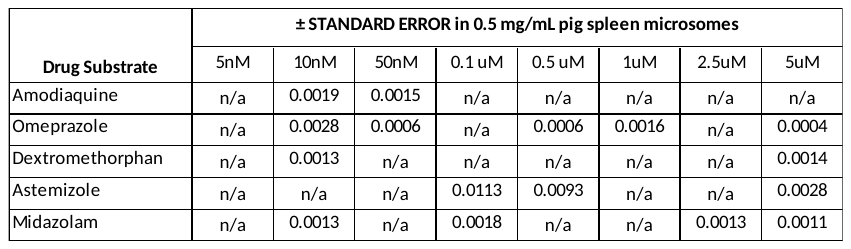

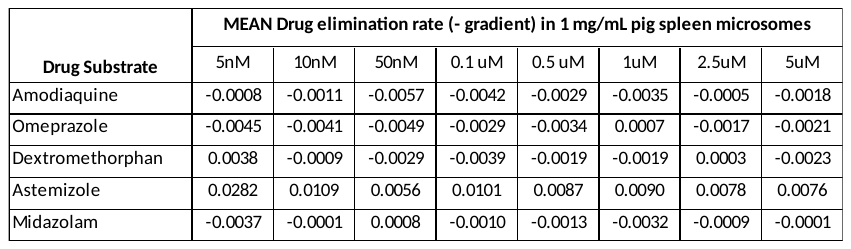

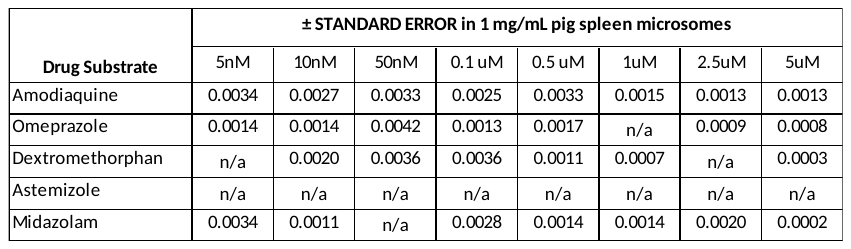

**Table. 3.0	UGT-mediated calculated Intrinsic clearance of 3 drug substrates in pig liver microsomes**

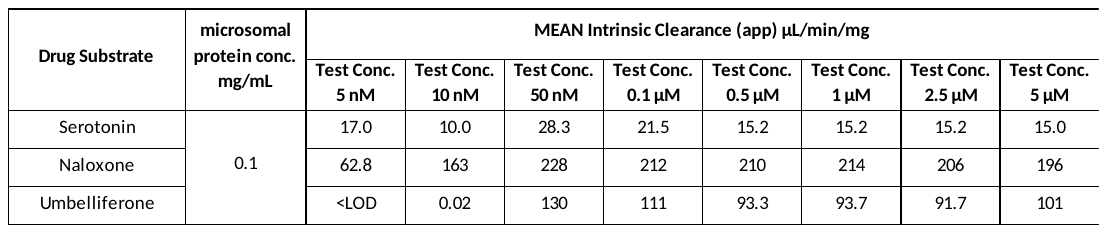

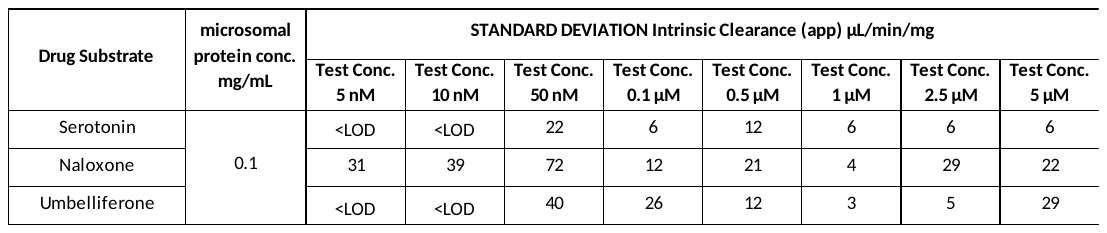

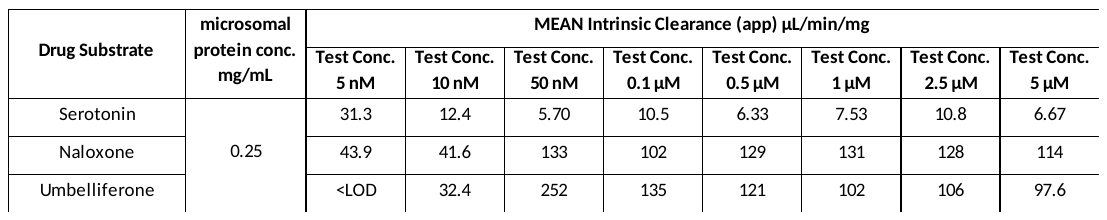

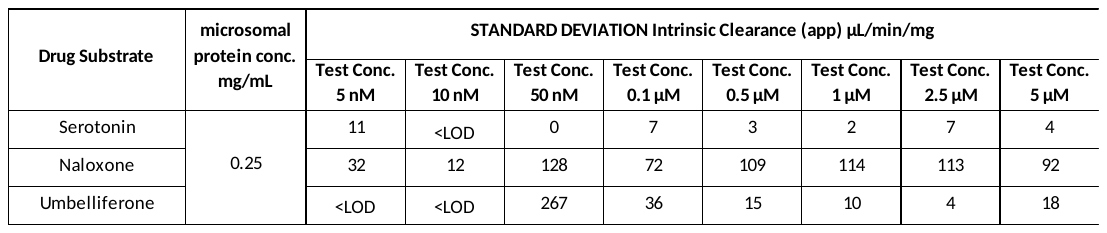

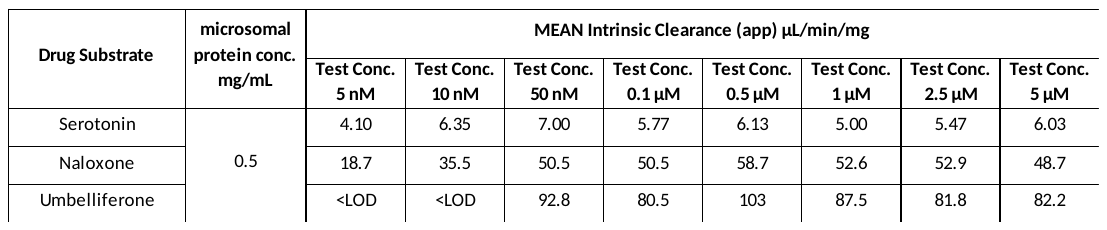

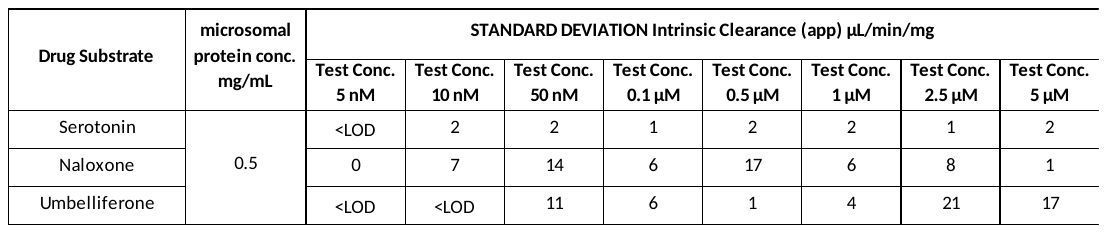

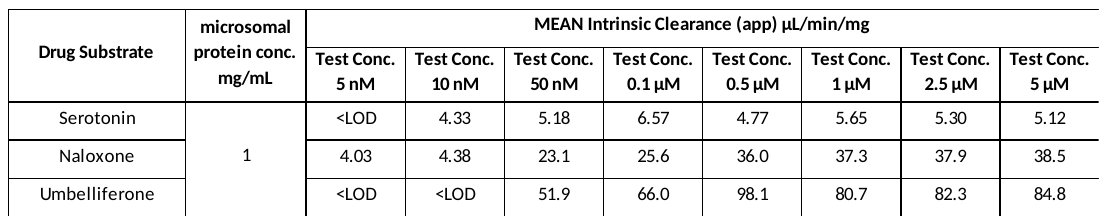

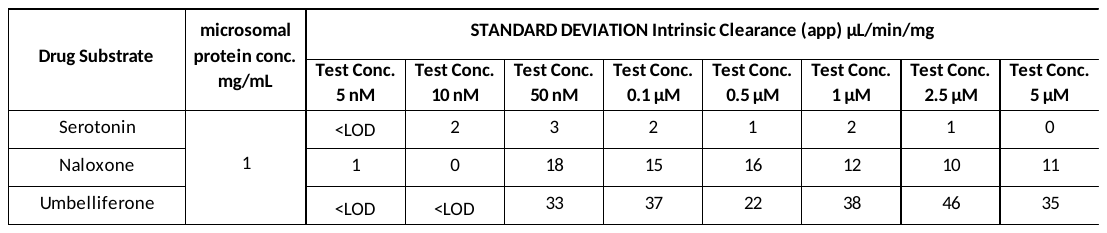

**Table. 3.1	MEAN UGT-mediated drug elimination rate** $\boldsymbol{k}$ **(- gradient) and ± standard error for 3 drug substrates in pig liver microsomes**

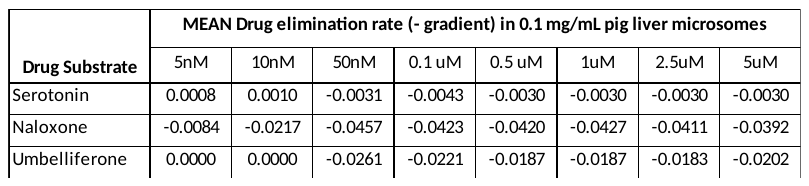

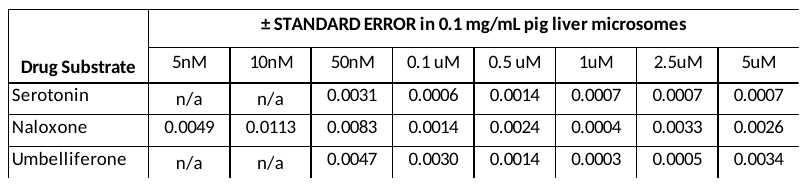

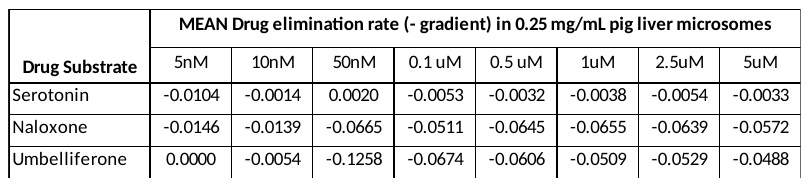

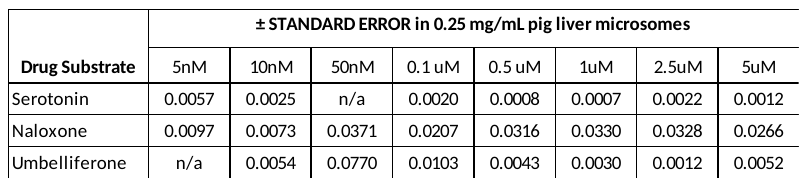

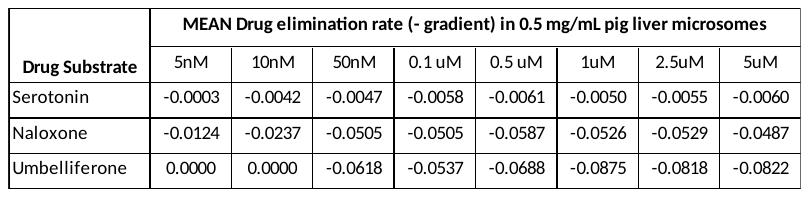

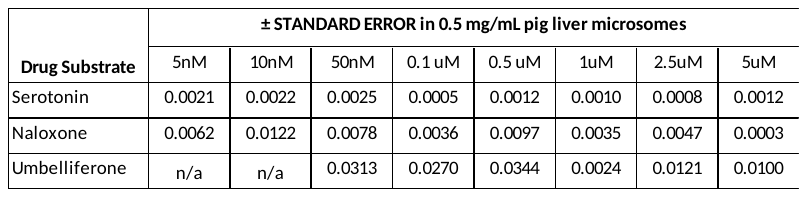

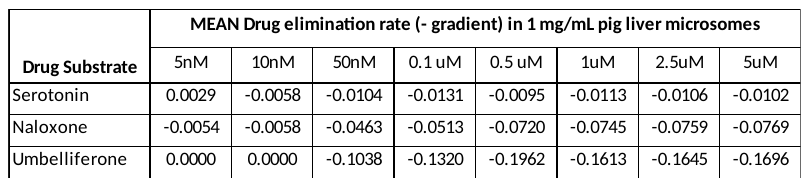

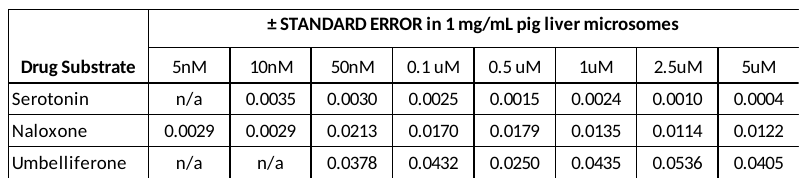

**Table. 4.0	UGT-mediated calculated Intrinsic clearance including ± standard deviation of 3 drug substrates in pig spleen microsomes**

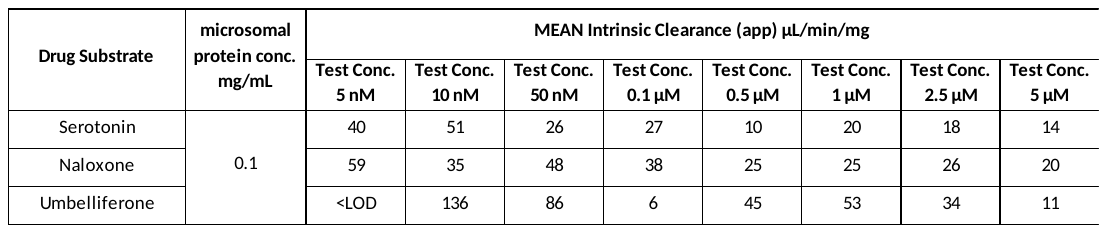

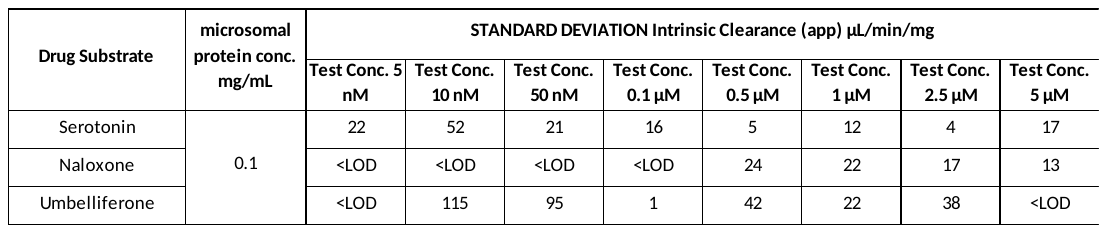

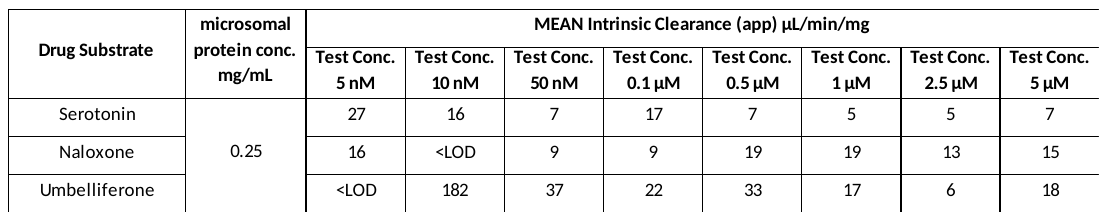

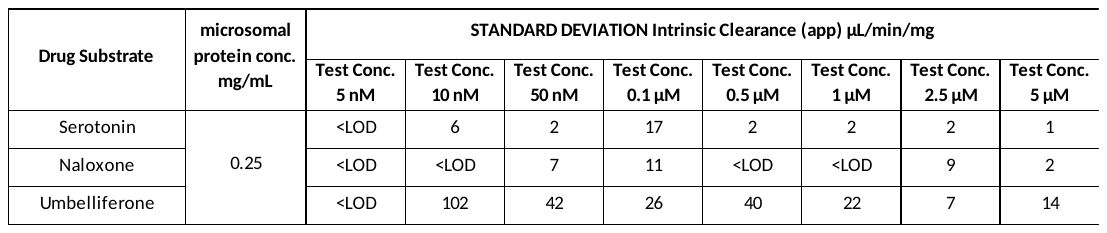

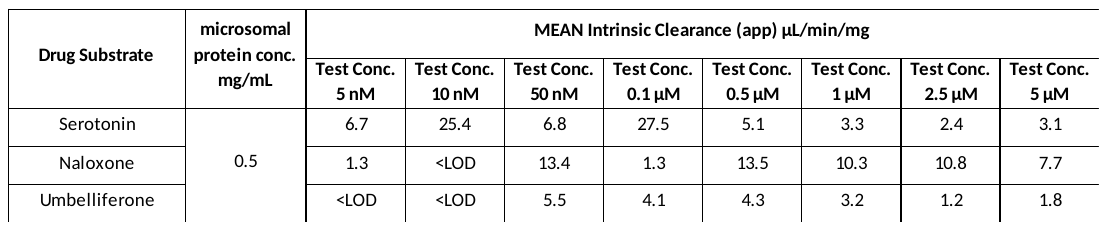

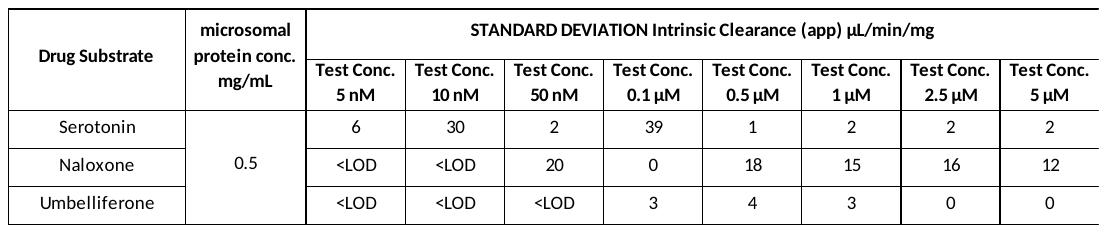

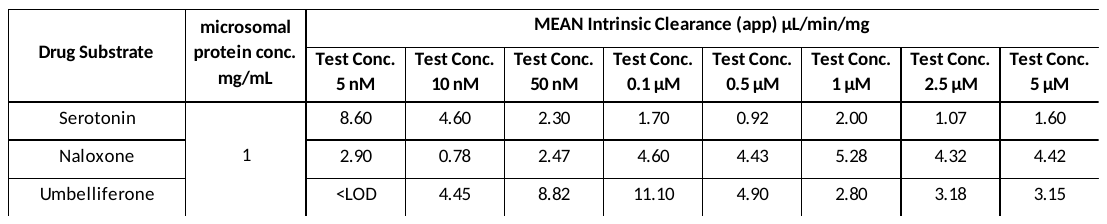

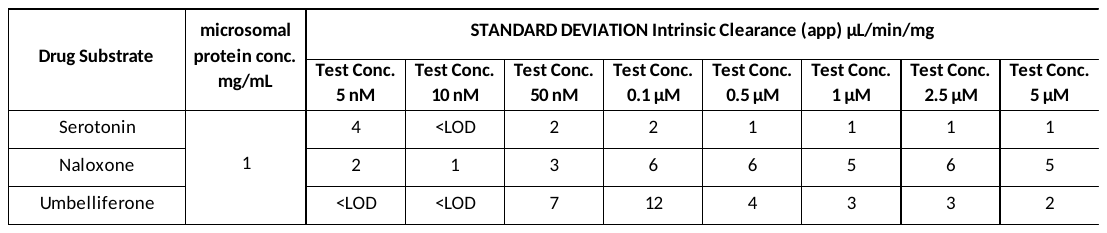

**Table. 4.1	MEAN UGT-mediated drug elimination rate** $\boldsymbol{k}$ **(- gradient) and ± standard error for 3 drug substrates in pig spleen microsomes**

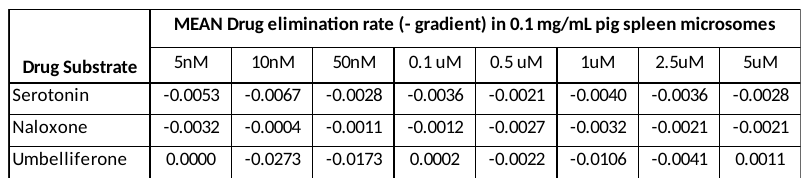

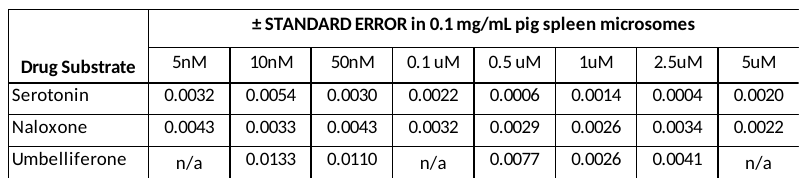

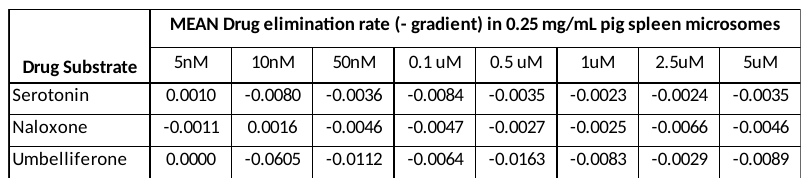

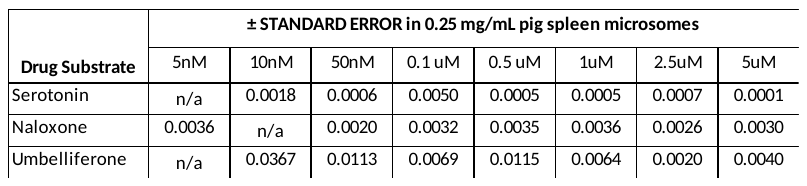

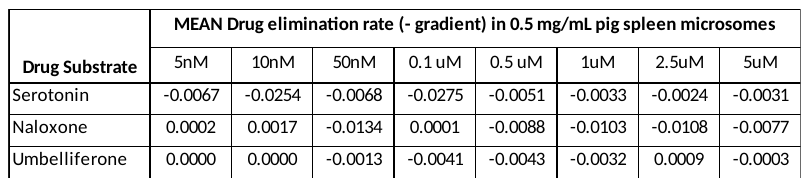

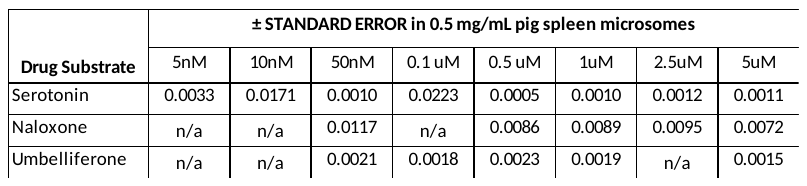

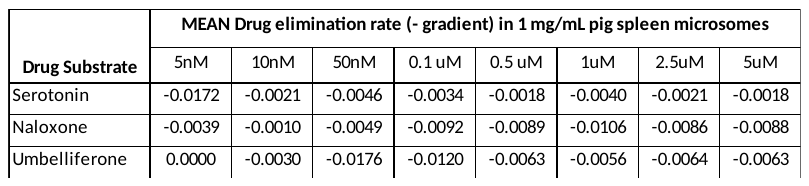

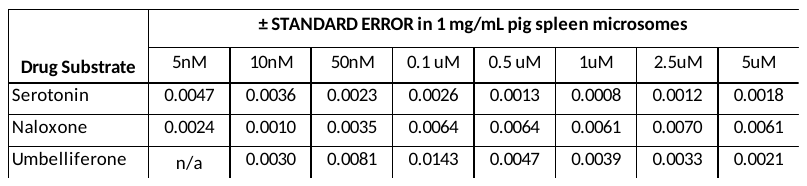

**Table. 5.0	CYP450-mediated calculated Intrinsic clearance including ± standard deviation of 5 drug substrates in pig hepatocytes**

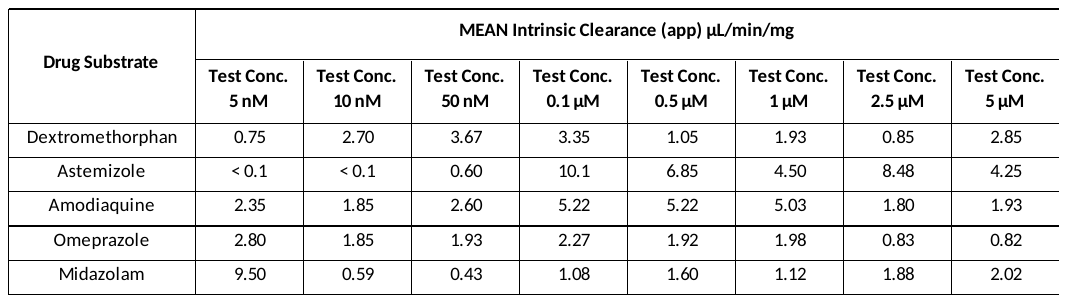

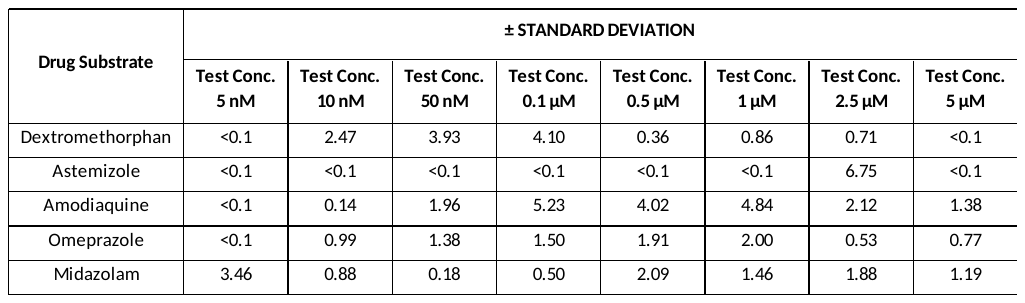

**Table. 5.1	MEAN CYP450-mediated drug elimination rate** $\boldsymbol{k}$ **(- gradient) and ± standard error for 5 drug substrates in pig hepatocytes**

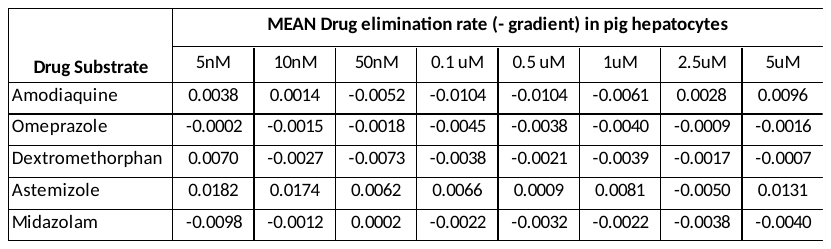

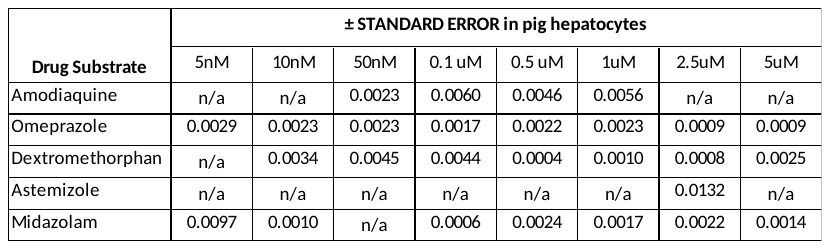

**Table. 6.0	CYP450-mediated calculated Intrinsic clearance including ± standard deviation of 5 drug substrates in pig splenocytes**

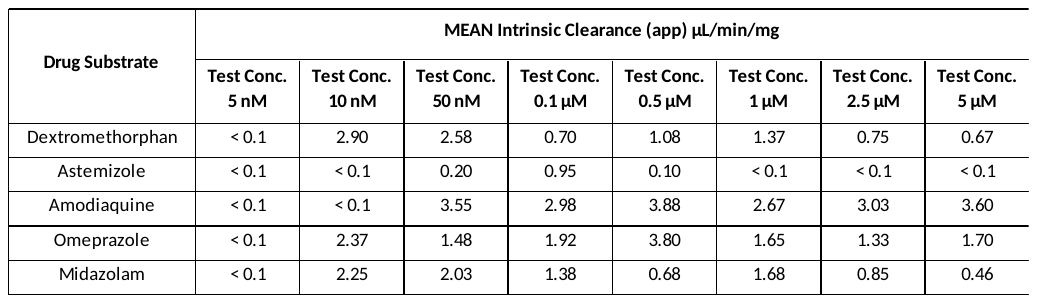

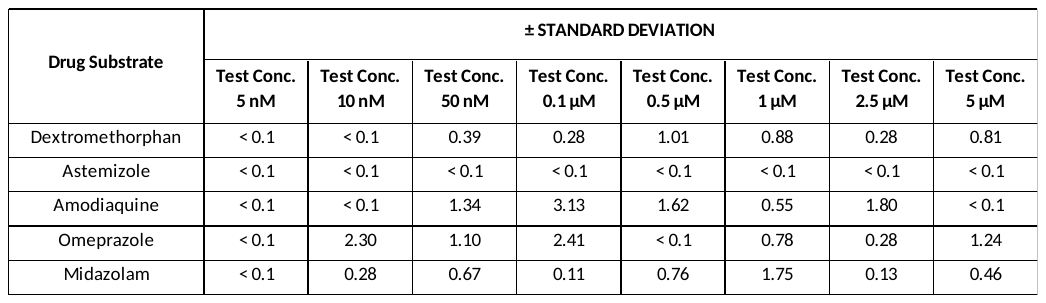

**Table. 6.1	MEAN CYP450-mediated drug elimination rate** $\boldsymbol{k}$ **(- gradient) and ± standard error for 5 drug substrates in pig splenocytes**

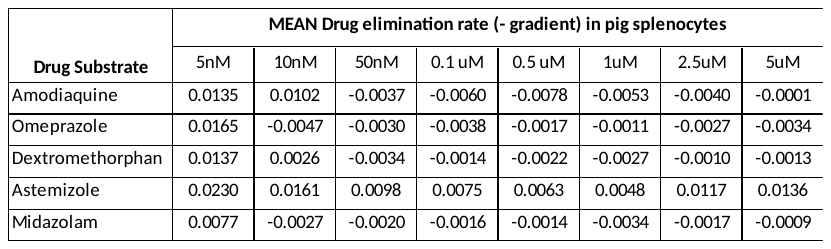

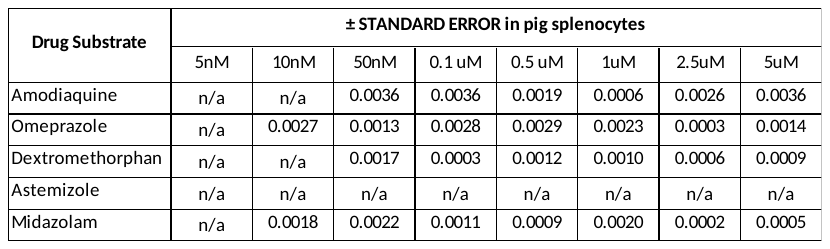

**Table. 7.0	CYP450-mediated MEAN metabolite formation levels n=3 (peak area response RAW data) monitored in pig liver microsomes, including ± standard deviation**

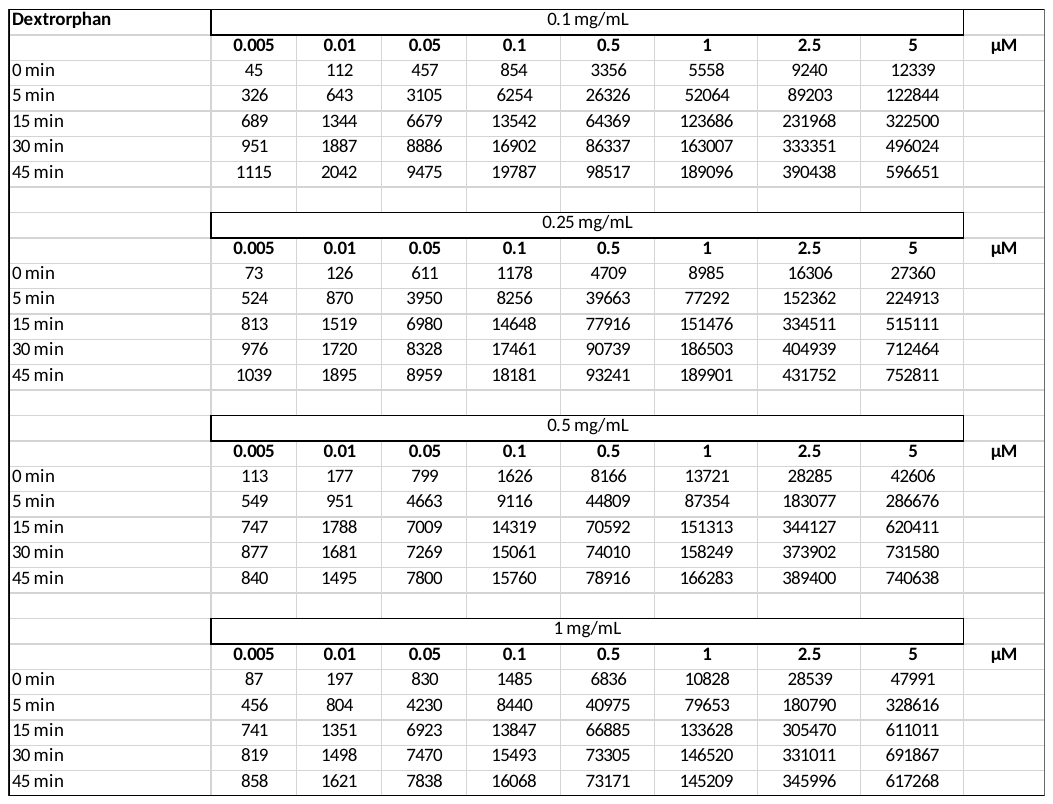

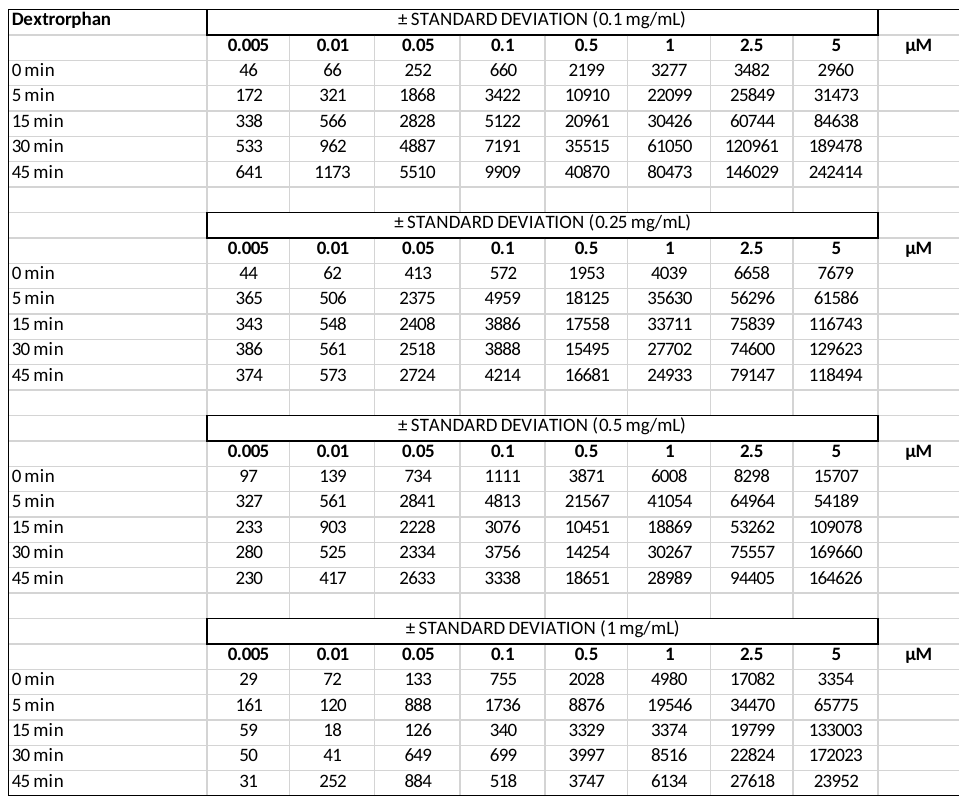

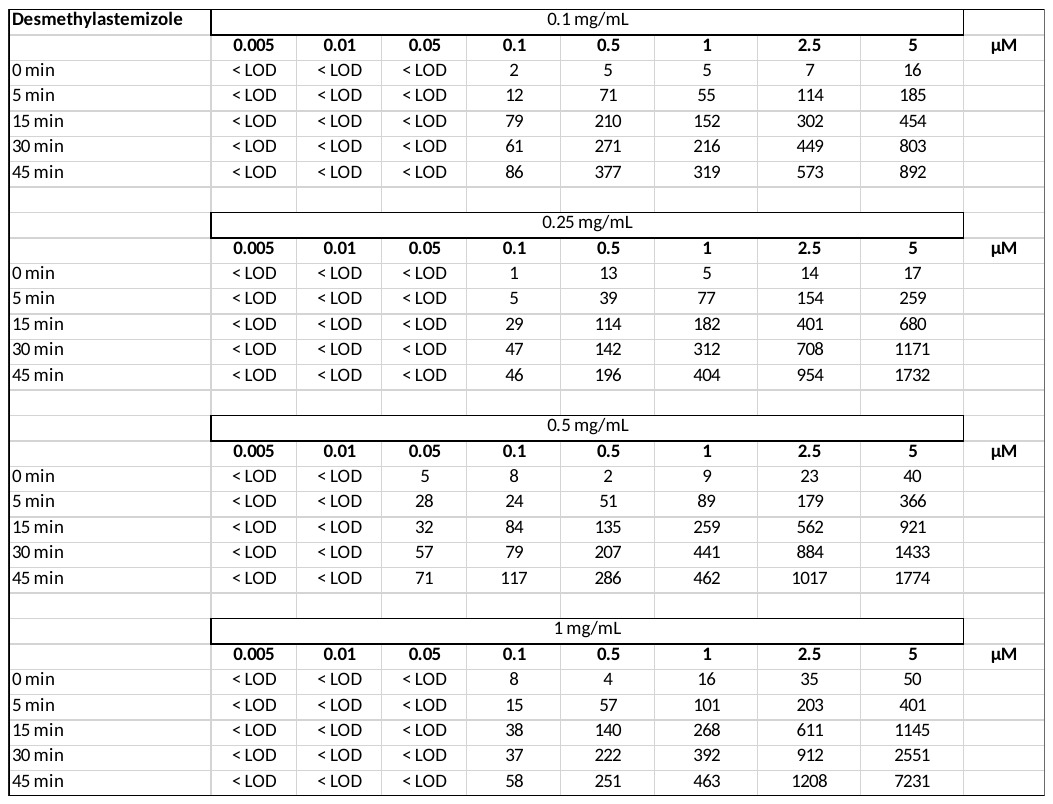

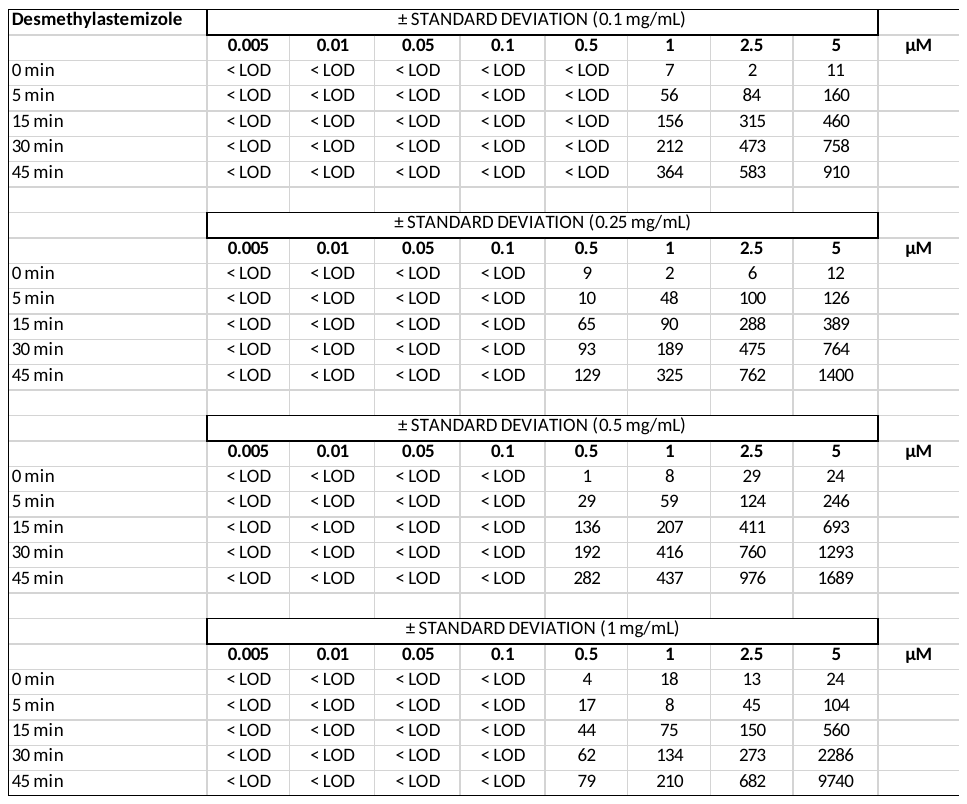

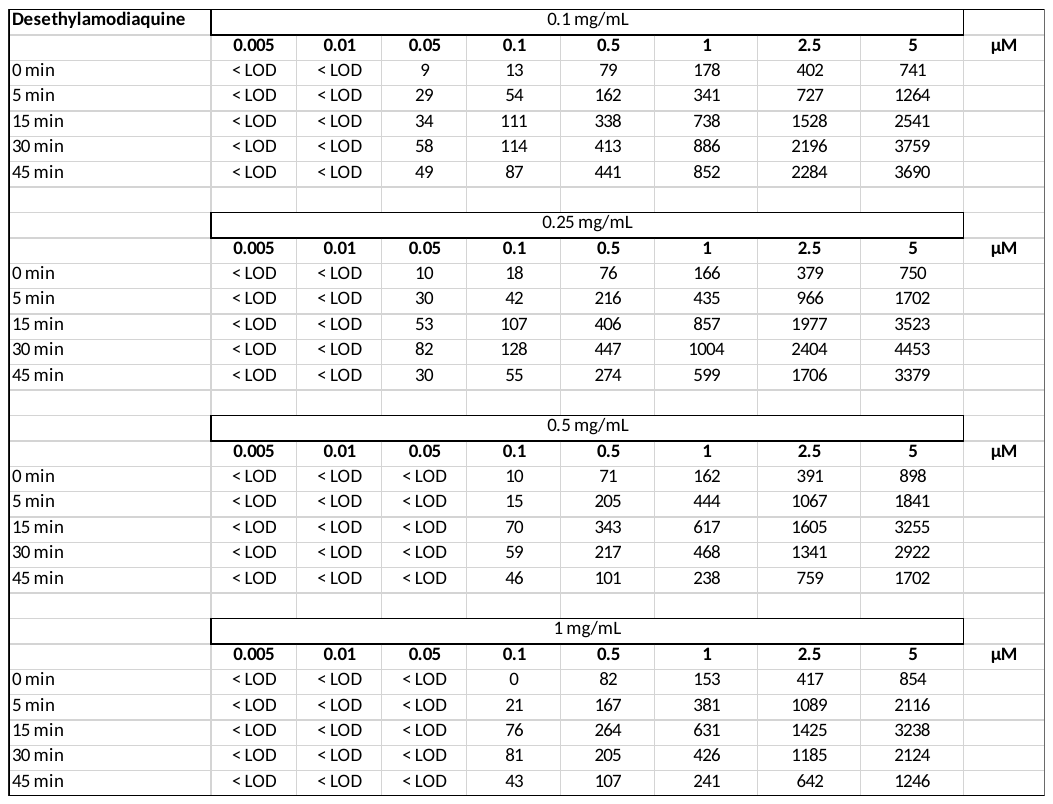

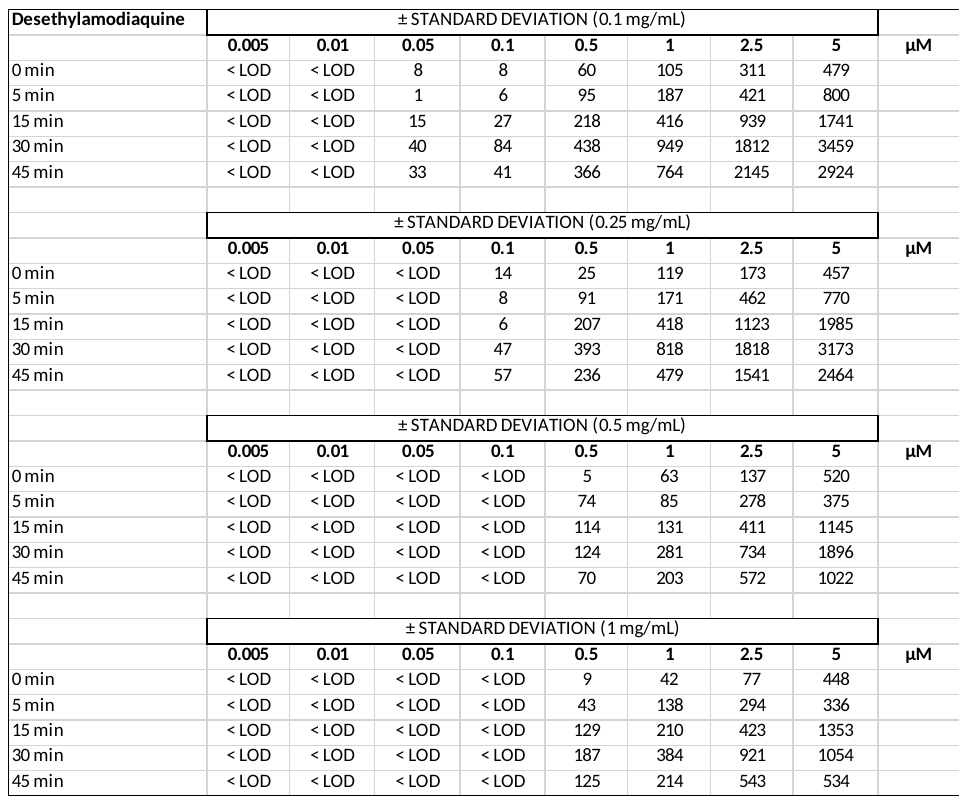

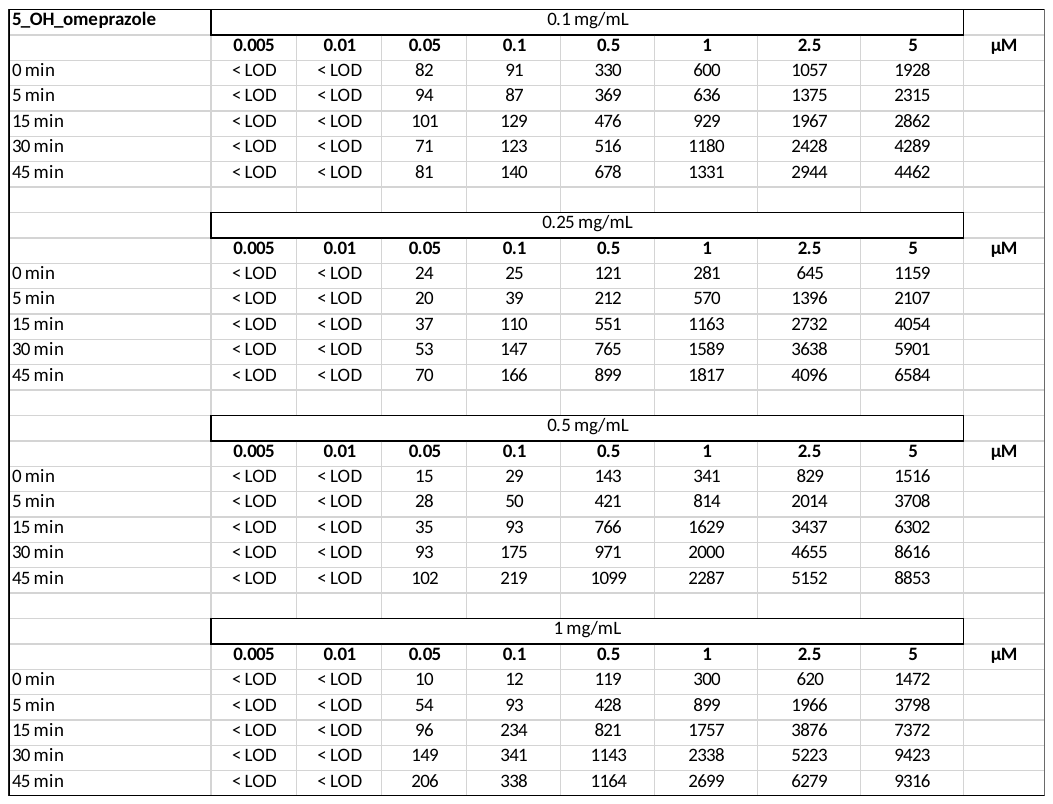

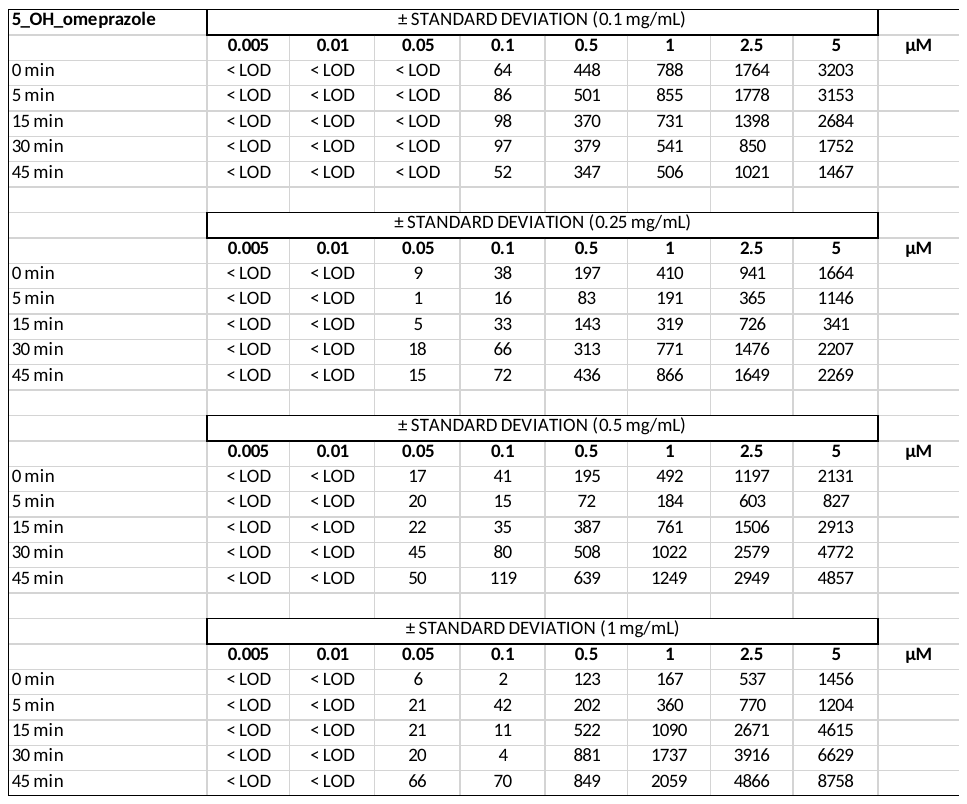

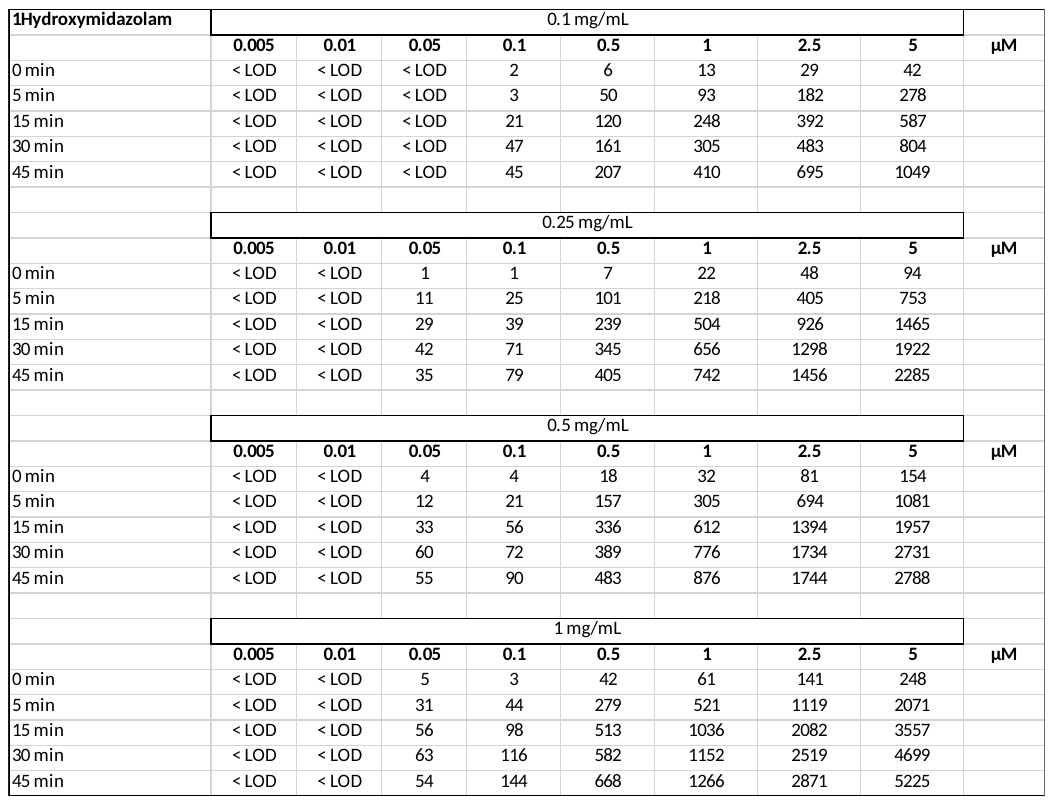

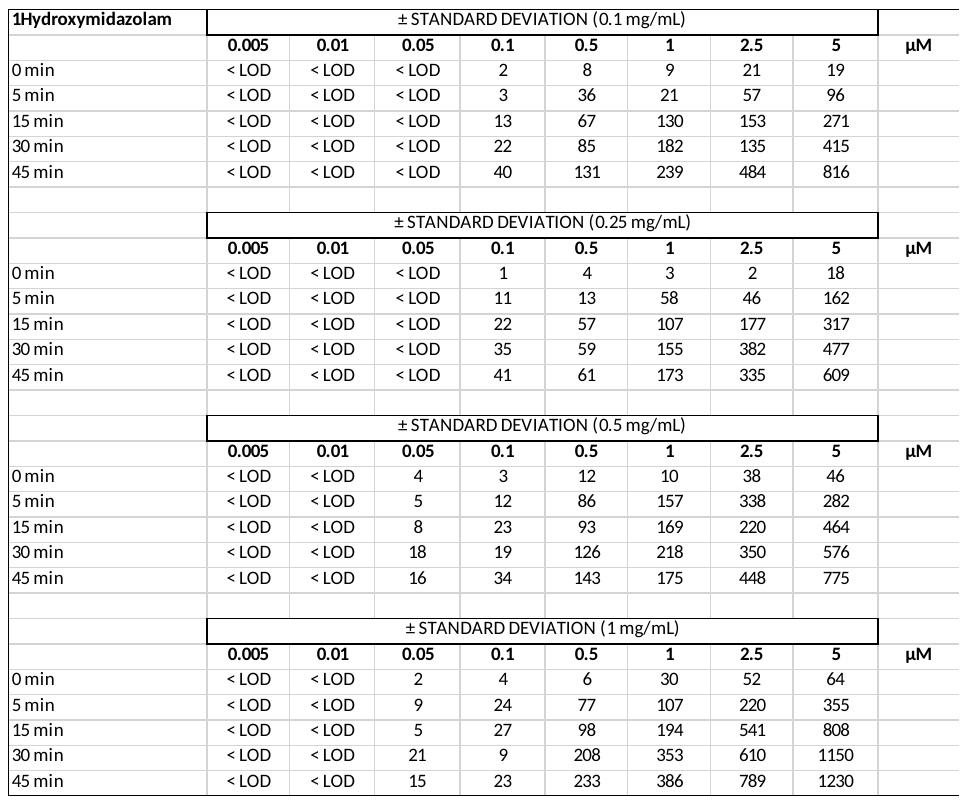

**Table. 8.0	CYP450-mediated MEAN metabolite formation levels n=3 (peak area response RAW data) monitored in pig spleen microsomes, including ± standard deviation**

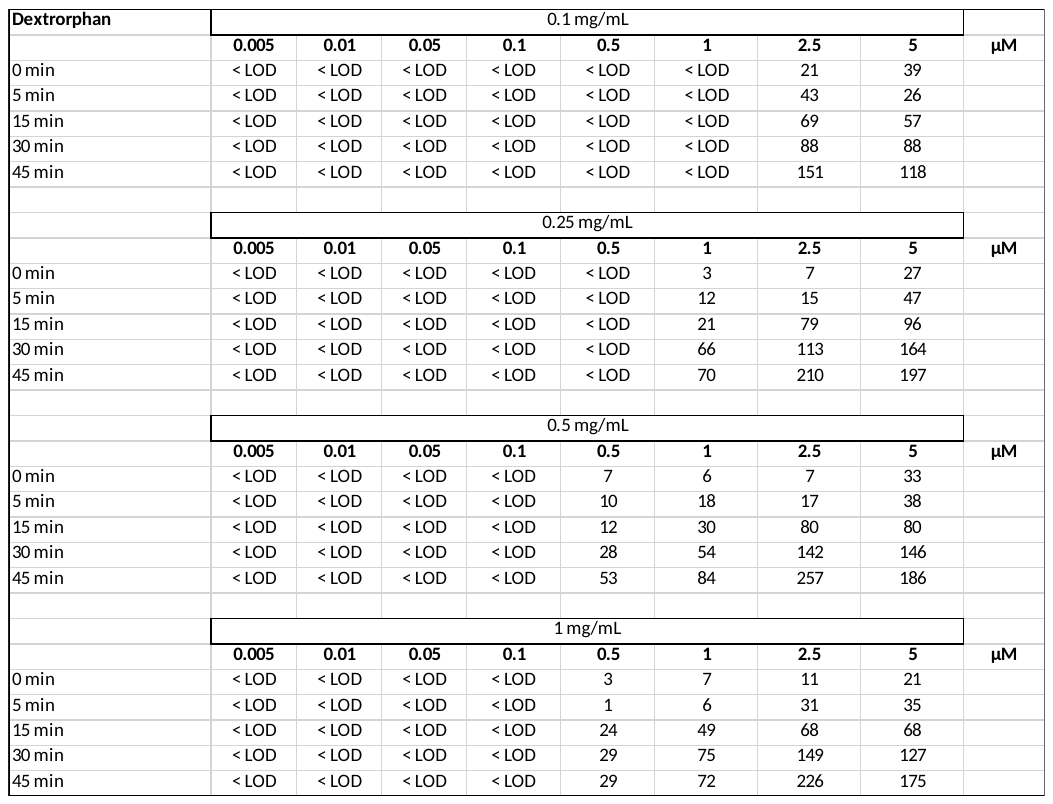

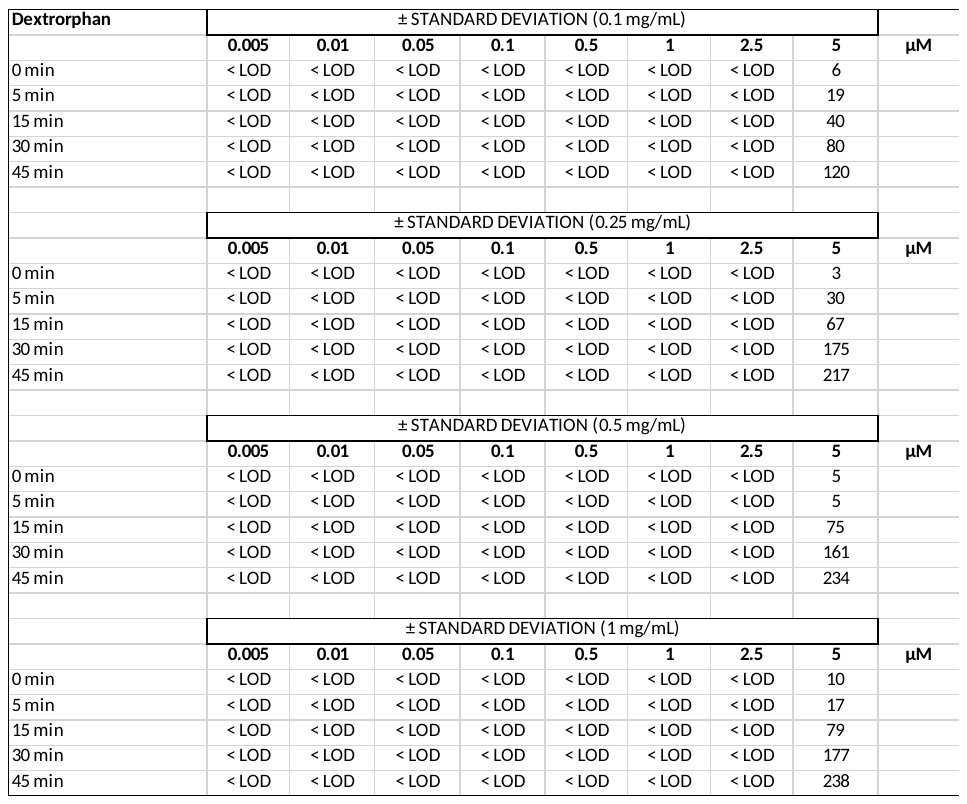

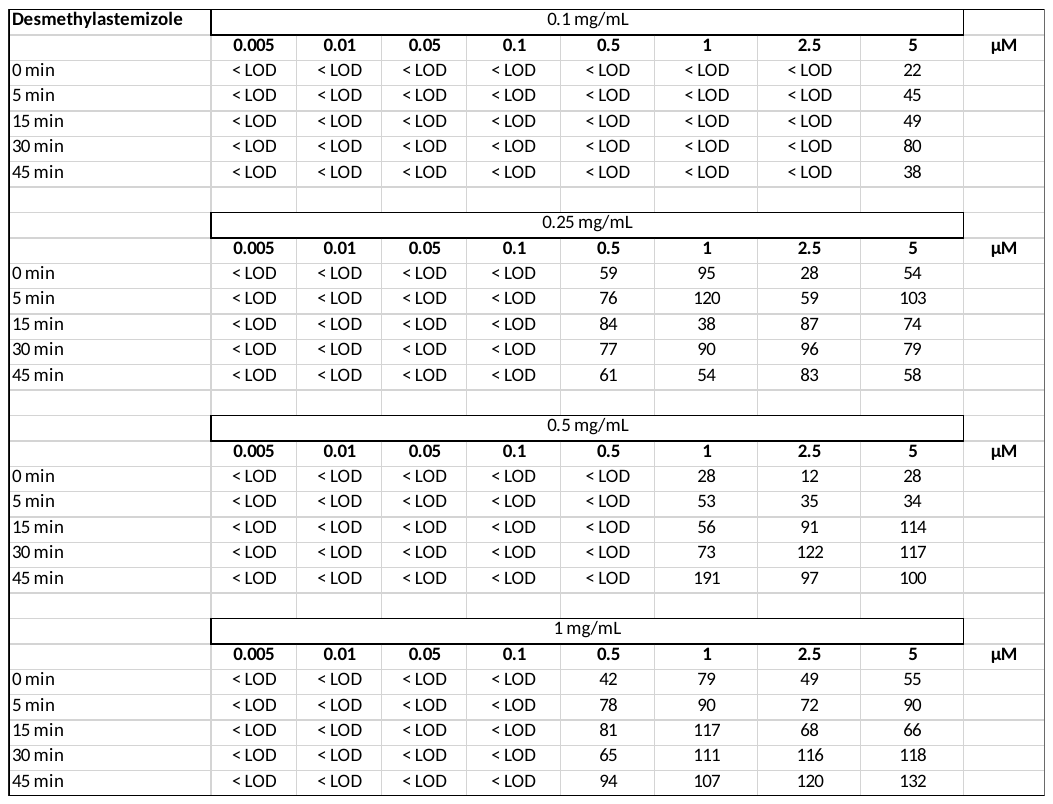

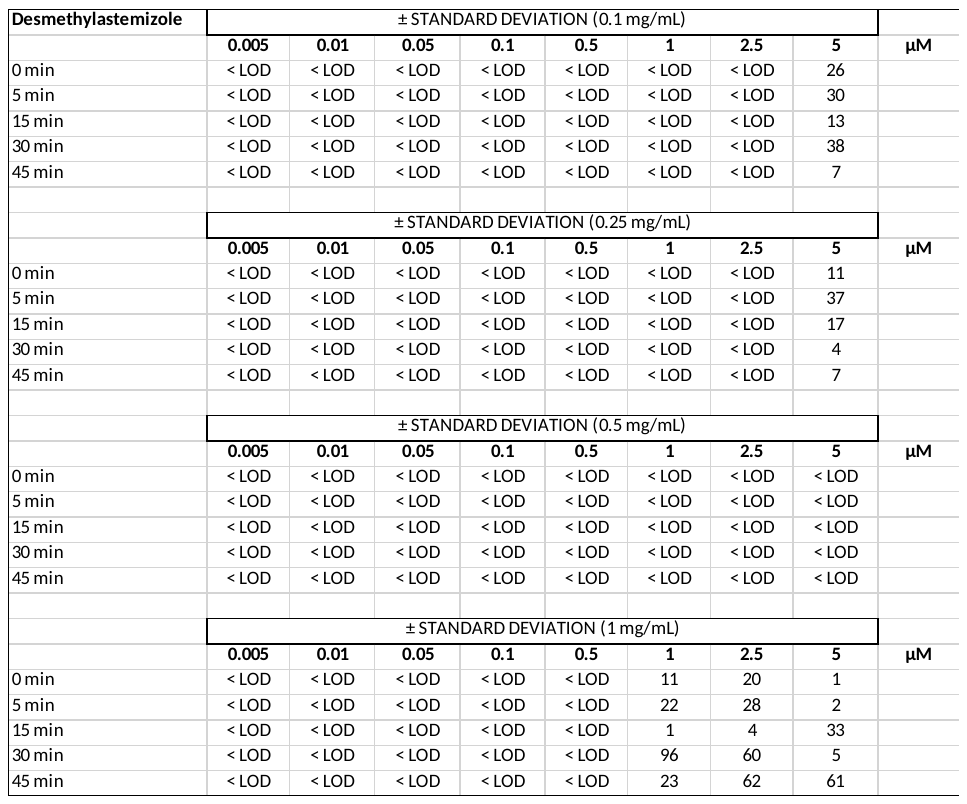

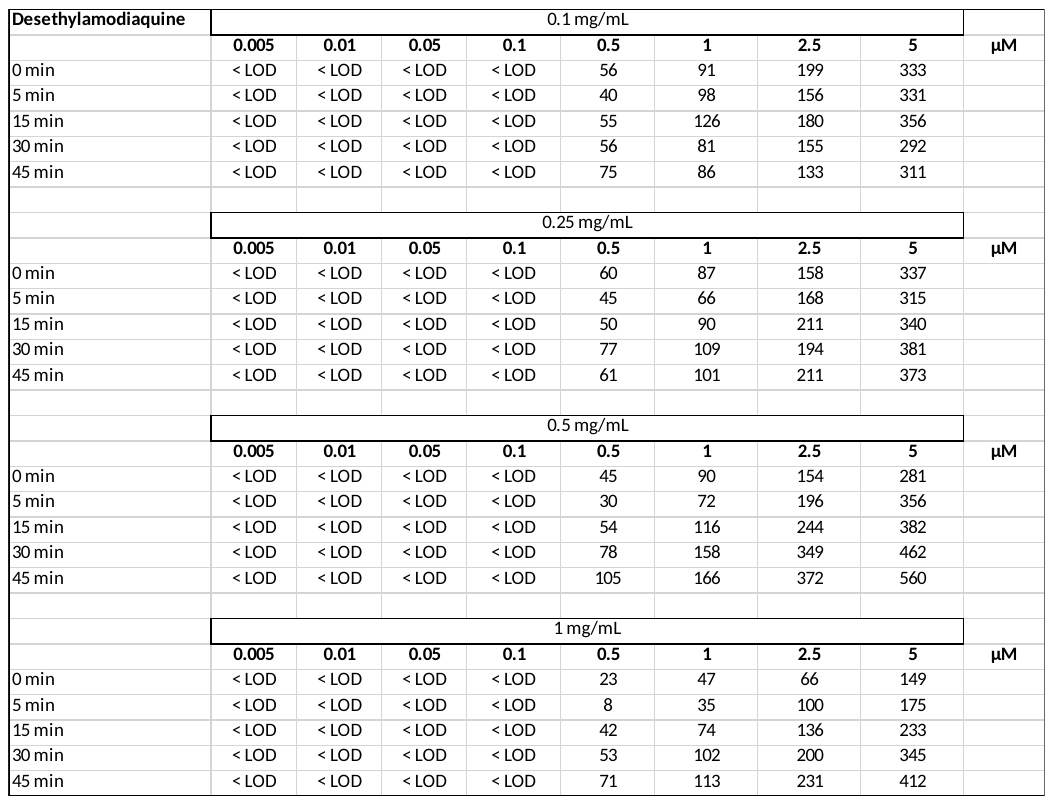

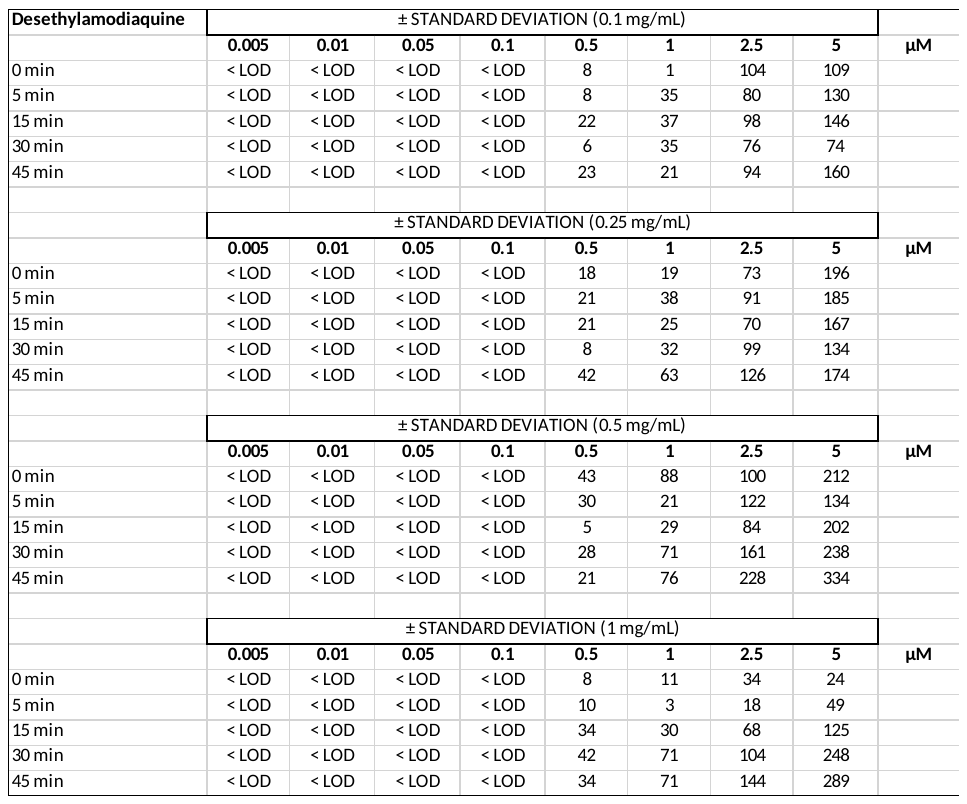

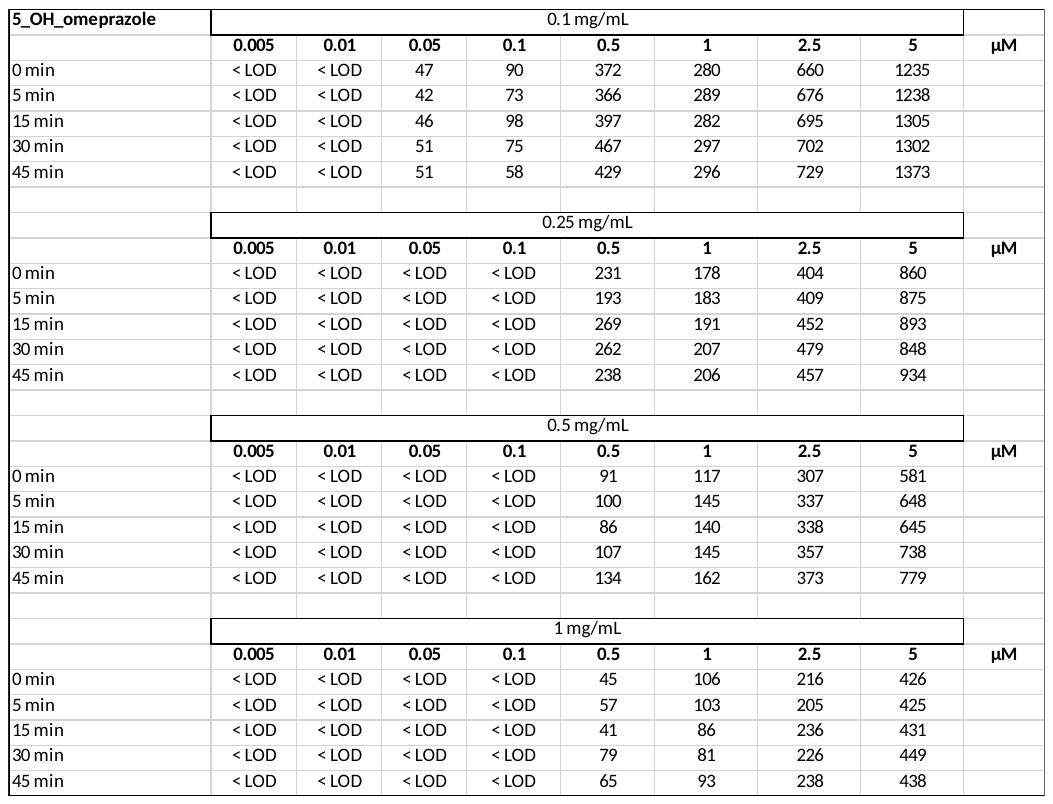

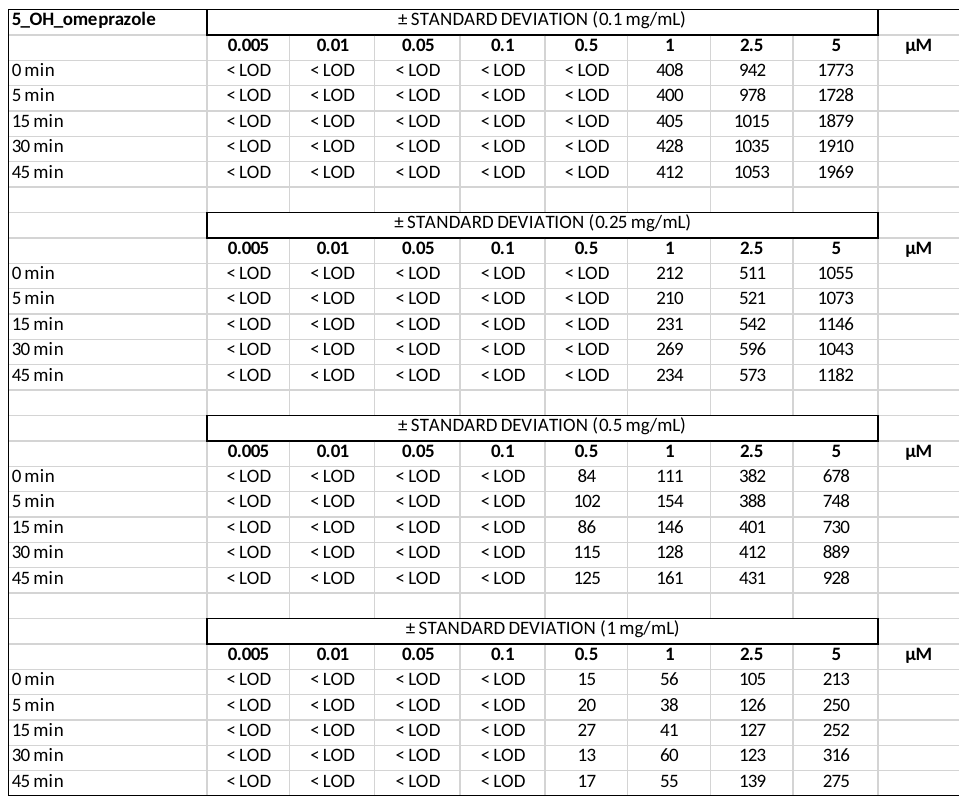

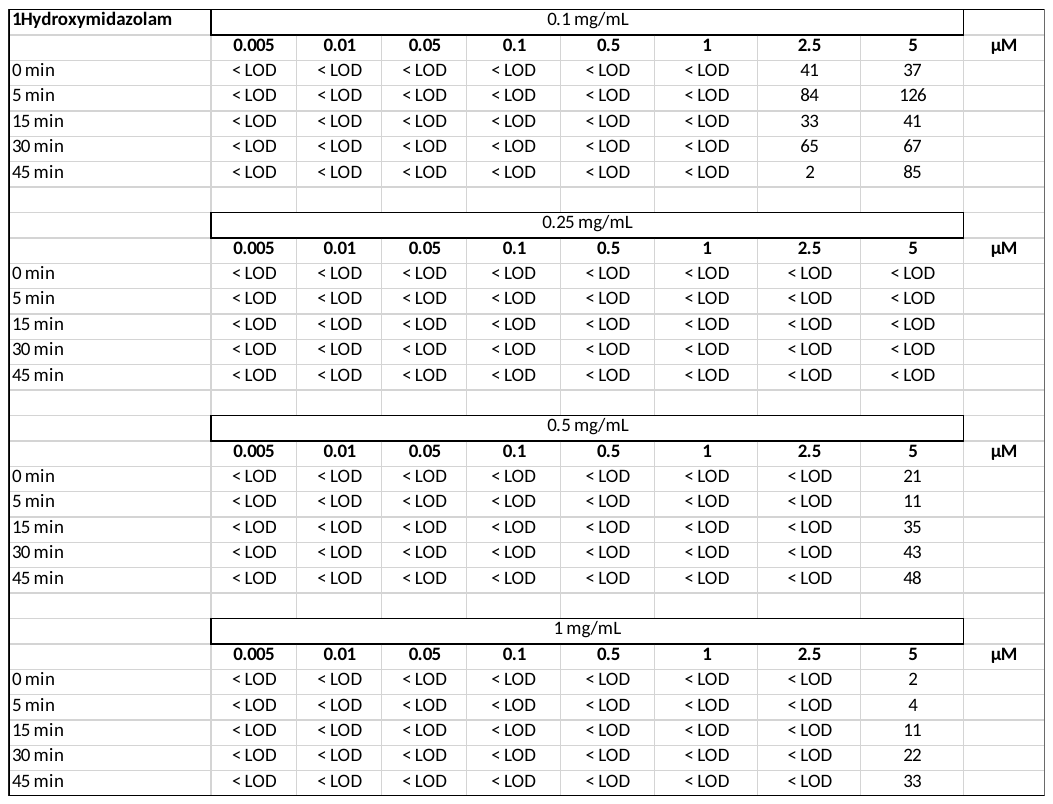

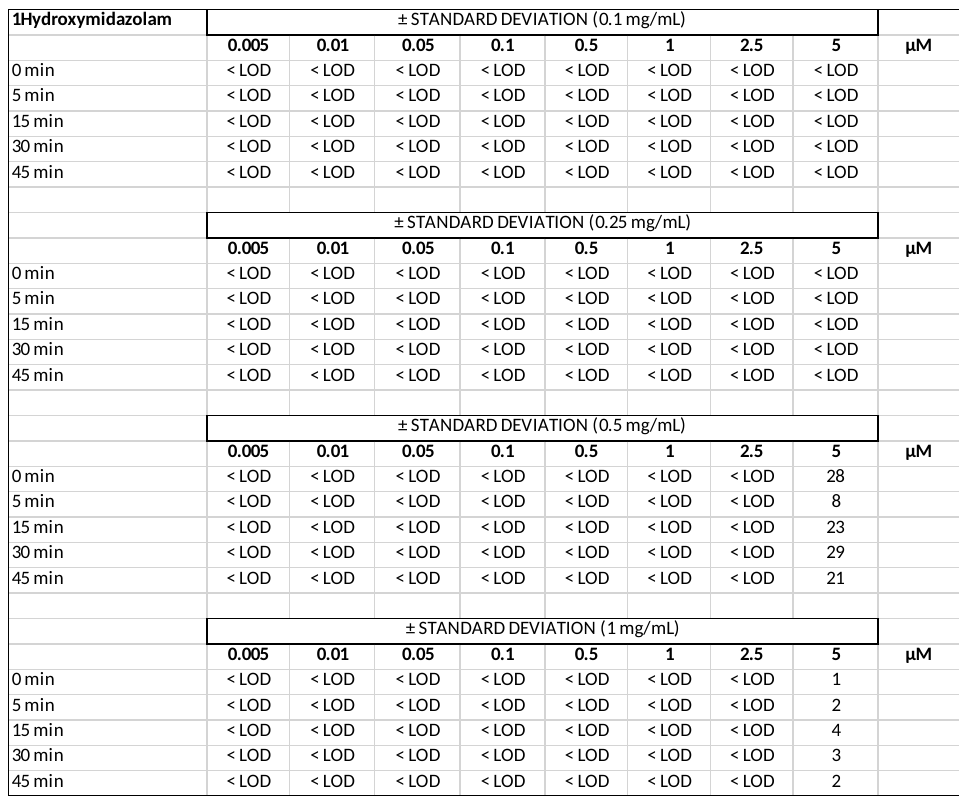

**Table. 9.0	UGT-mediated MEAN metabolite formation levels n=3 (peak area response RAW data) monitored in pig liver microsomes, including ± standard deviation**

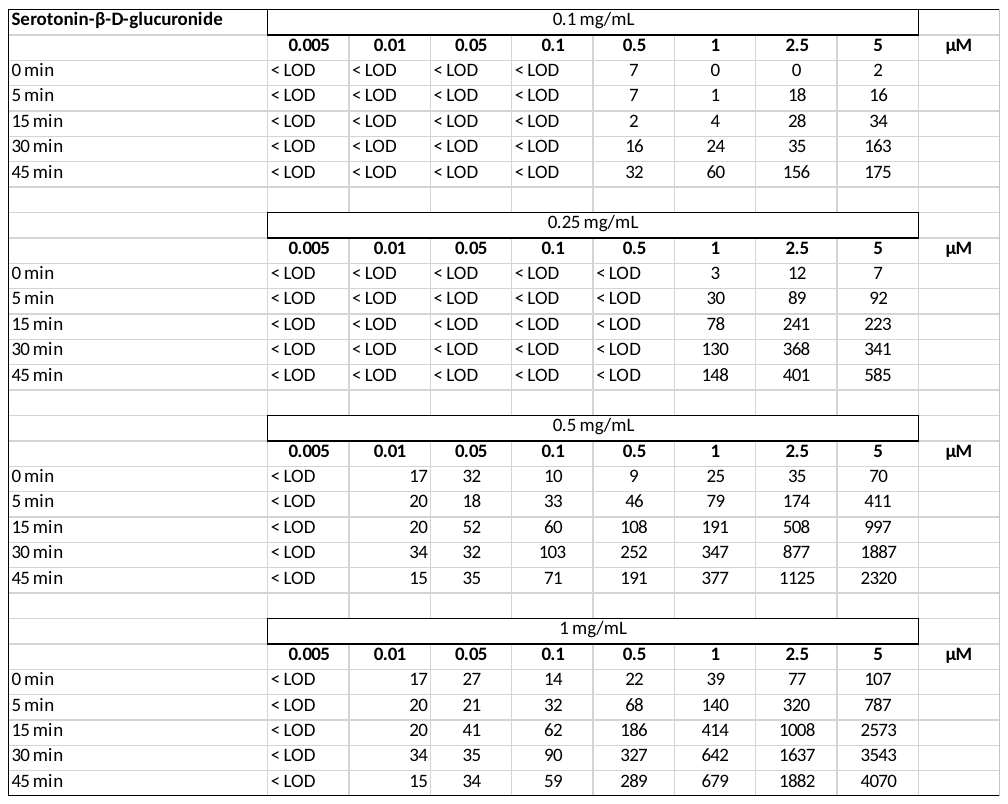

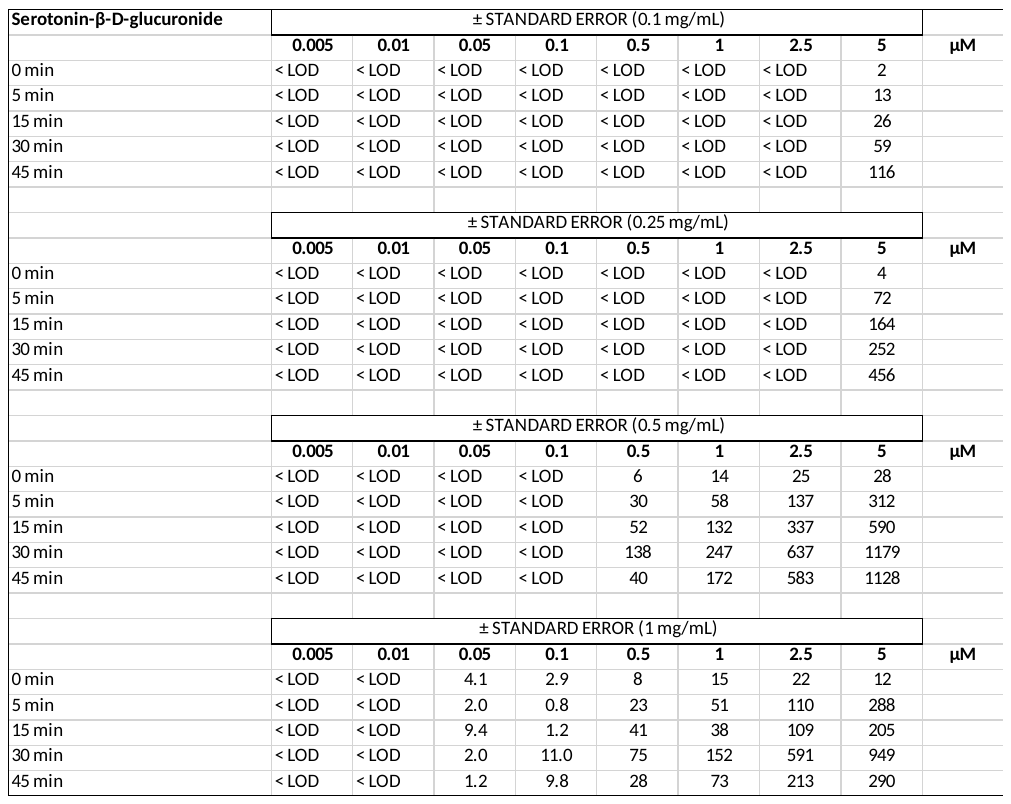

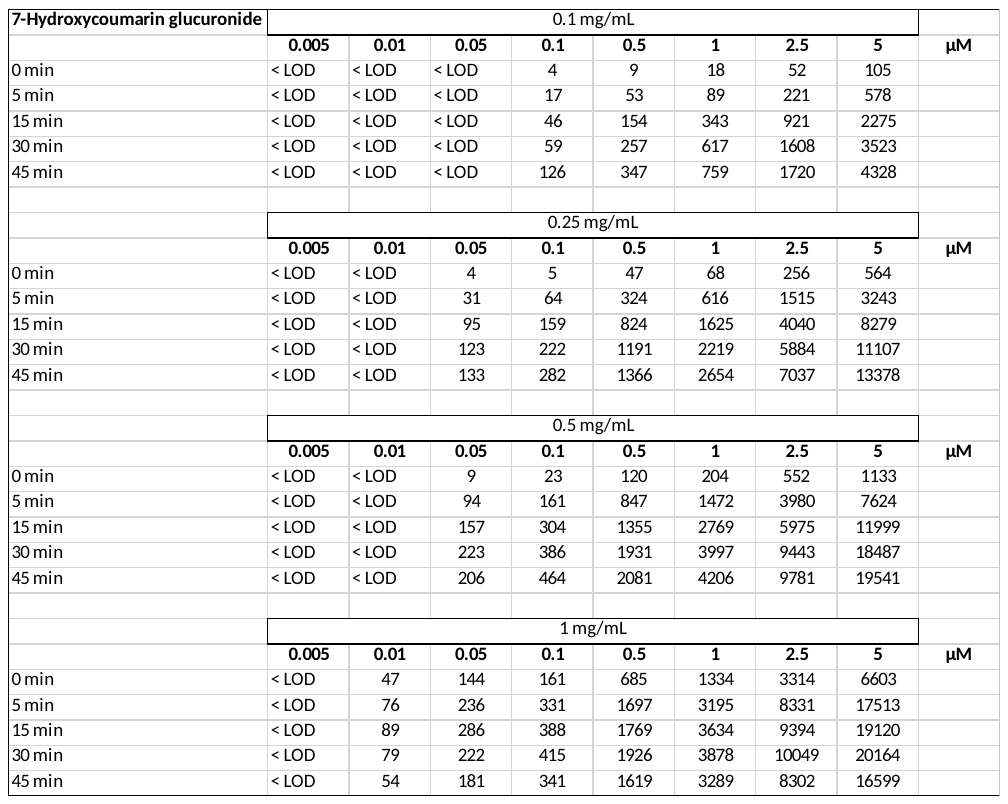

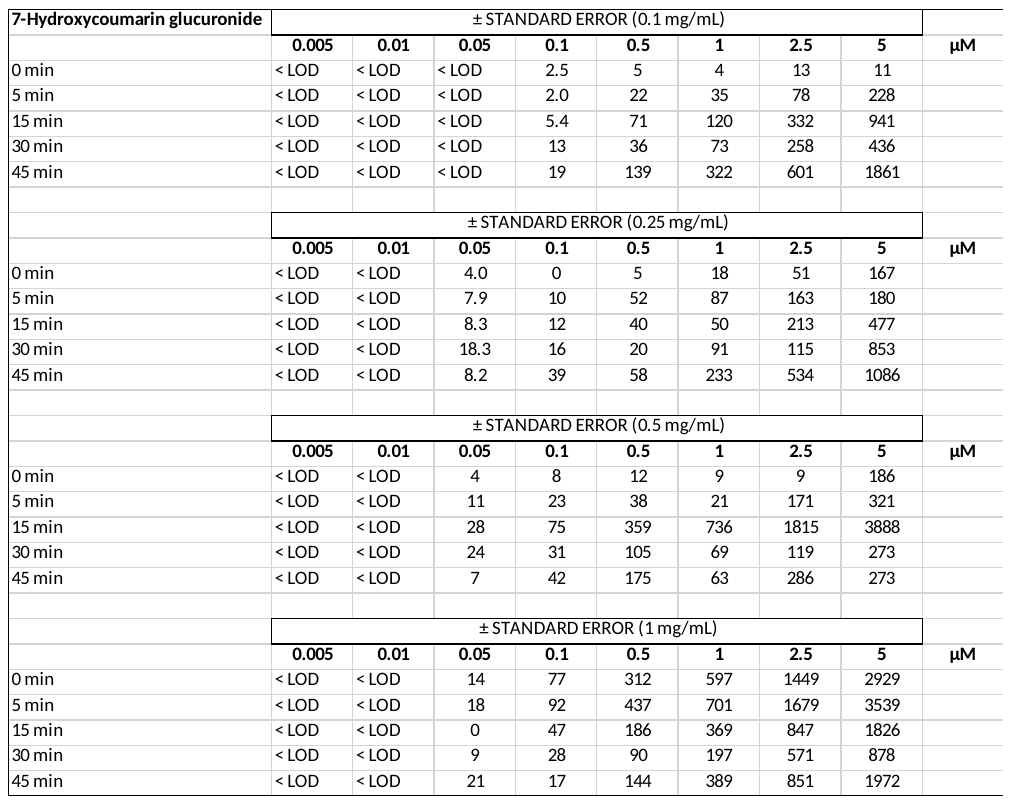

**Table. 10.0	UGT-mediated MEAN metabolite formation levels n=3 (peak area response RAW data) monitored in pig spleen microsomes, including ± standard deviation**

**Table. 11.0	CYP450-mediated MEAN metabolite formation levels n=3 (peak area response RAW data) monitored in pig hepatocytes, including ± standard deviation**

**Table. 12.0	CYP450-mediated MEAN metabolite formation levels n=3 (peak area response RAW data) monitored in pig splenocytes, including ± standard deviation**
